# Tempo and mode of evolution across multiple traits in an adaptive radiation of birds (Vangidae)

**DOI:** 10.1101/2024.12.10.627847

**Authors:** Anya Lilith Becker Auerbach, Euan Horng Jiunn Lim, Sushma Reddy

## Abstract

An ongoing challenge in macroevolutionary research is identifying common drivers of diversification amid the complex interplay of many potentially relevant traits, ecological contexts, and intrinsic characteristics of clades. In this study, we used geometric morphometric and phylogenetic comparative methods to evaluate the tempo and mode of trait evolution in the adaptive radiation of Malagasy vangas and their mainland relatives. The Malagasy radiation is more diverse in both skull and foot shape. However, rather than following the classic “early burst” of diversification, trait evolution accelerated well after their arrival in Madagascar, likely driven by the evolution of new modes of foraging and especially of a few species with highly divergent morphologies. Each anatomical region showed differing evolutionary patterns, and the presence of morphological outliers impacted the results of some analyses, particularly of integration and modularity. Our results demonstrate that the adaptive radiation of Malagasy vangas has evolved exceptional ecomorphological diversity along multiple, independent trait axes, mainly driven by a late expansion in niche space due to key innovations. Our findings highlight the evolution of extreme forms as an overlooked feature of adaptive radiation warranting further study.

## Introduction

Adaptive radiations – clades which have undergone an exceptional degree of ecological diversification – are emblematic examples of evolution and offer powerful opportunities to help understand the processes that generate ecological and phenotypic diversity (Givnish and Sytsma 1997; Schluter 2000; Hodges and Derieg 2009). Despite substantial interest in the study of adaptive radiation, major disagreements remain regarding 1) how these clades should be identified, and 2) the generalizability and predictive power of these patterns as evolutionary models (Losos and Miles 2002; Gavrilets and Losos 2009; Olson and Arroyo-Santos 2009; Moen et al. 2021). Many shared general patterns have been proposed for adaptive radiations, but most are not generalizable across the diversity of clades usually considered under this framework, making identification of radiations and their comparative study difficult (Gillespie et al. 2020). Also poorly understood is the degree to which adaptive radiation is predictable, in terms of both the external factors and intrinsic features of clades which may promote or inhibit radiation (Yang 2001; Lovette et al. 2002a; Kassen 2009; Glor 2010; Losos and Mahler 2010; Wellborn and Langerhans 2015; Stroud and Losos 2016). Although many studies have examined rates of evolutionary change in radiations, understanding their predictability will require reconciling variable patterns of diversification across multiple traits which may be subject to different selective pressures.

One of the most common patterns used to identify adaptive radiations is an “early burst” – but an overemphasis on this criterion risks obscuring true complexity by excluding other modes of diversification. The early burst mode has often been considered a defining feature because classically, adaptive radiations are predicted to undergo rapid early diversification in response to some ecological opportunity, followed by declining rates of diversification as niche space is filled (Simpson 1944; Schluter 2000; Freckleton and Harvey 2006; Gavrilets and Losos 2009; Martin and Richards 2019). Until recently, the emphasis on testing for an early burst resulted in a focus on speciation rates, while the quantification of ecological diversity was often overlooked (Yoder et al. 2010; Givnish 2015). More recently, methodological advances have enabled a shift towards testing for an early burst using trait data and other means of quantifying ecological diversity (Harmon et al. 2010; Mahler et al. 2010; Slater et al. 2010). These studies have revealed that, while rapid speciation appears to be correlated with ecological trait diversification on average (Rabosky et al. 2013; Cooney and Thomas 2021), these processes are frequently decoupled in individual clades (Derryberry et al. 2011; Venditti et al. 2011; Folk et al. 2019; Martin and Richards 2019; Rowsey et al. 2019; Barreto et al. 2023). Furthermore, the framework of comparing support for a limited set of diversification scenarios (e.g., early burst vs Brownian motion) can lead to misidentification of other, often more complex models (Martin et al. 2023). Although an early burst can be a strong indicator of diversification in response to ecological opportunity, defining adaptive radiation solely by this criterion has the potential to limit our understanding of how different conditions may impact the tempo of diversification (Losos and Mahler 2010; Astudillo-Clavijo et al. 2015).

Identifying appropriate ecologically relevant traits to target for a given clade is itself an ongoing challenge. Many studies focus on single, univariate measurements such as body size, despite conceptual models of adaptive radiation which often involve simultaneous and/or sequential divergence in multiple aspects of adaptive changes (Slater 2022). The results from any single trait risk being fundamentally misleading if diversification has occurred along multiple trait axes. The appropriate selection of traits for a particular clade, and how to consider simultaneous or sequential diversification of multiple traits, are underexplored in the literature (Ackerly et al. 2006; Donoghue and Sanderson 2015; Slater and Friscia 2019; Mutumi et al. 2023). Examining a variety of traits will best represent the underlying ecological diversity present in different clades, and this attention to organismal biology and natural history knowledge remains essential for drawing robust conclusions (Losos 2010). Different ecologically relevant traits may show different patterns of diversification in the same clade, as in the Malagasy radiation of mantellid frogs, where evolutionary rates of shape but not size or performance-related metrics were elevated relative to other anuran clades (Moen et al. 2021). We should expect adaptive radiation to frequently proceed along multiple axes of diversification, and in these cases understanding the evolutionary dynamics of these traits in concert is central to understanding how diversity is generated.

Another major area of evolutionary research concerns understanding variation in the intrinsic capacity of a clade to diversify, i.e. its evolvability. Several factors have been proposed to explain why, even when seemingly presented with similar ecological opportunity, some clades diversify so spectacularly whereas others do not (Lovette et al. 2002a; Sidlauskas 2008; Wellborn and Langerhans 2015; Jablonski 2022). A key characteristic impacting evolvability is the degree of independence between aspects of an organism’s phenotype (Kirschner and Gerhart 1998; Yang 2001; Jablonski 2022). Traits less able to vary independently of one another are described as more integrated, whereas modularity refers to the organization of traits into discrete “quasi-independent" regions (modules). Integration and modularity impact evolutionary rates and trajectories at multiple levels-genetic, developmental, and evolutionary (Cheverud 1984; Schluter 1996; Klingenberg 2008; Conith et al. 2021; Zelditch and Goswami 2021). We focus here on evolutionary integration as measured by comparing patterns of morphological trait covariation across a clade.

Theoretical and empirical studies have shown mixed evidence for how integration and modularity may shape adaptive radiations. Higher modularity has usually been thought to promote evolvability, with greater trait independence permitting a wider range of phenotypes to evolve (Wagner and Altenberg 1996; Yang 2001; Felice and Goswami 2018; Larouche et al. 2018; Walter et al. 2018; Burress et al. 2020). However, other studies suggest that integration can facilitate the evolution of more disparate phenotypes by maintaining key functional relationships and promoting rapid evolution along paths of least resistance (Schluter 1996; Griswold 2006; Goswami et al. 2014; Hedrick et al. 2020; Navalón et al. 2020; Evans et al. 2021). Continued work to uncover how integration and modularity shape rates and patterns of trait evolution is necessary to begin to understand the wide variation in evolvability we observe across clades (Felice et al. 2018).

The adaptive radiation of Malagasy vangas is an ideal system in which to explore these questions regarding the tempo and mode of diversification. The forty species in the family Vangidae are shrike-like birds found in tropical Asia and Africa (Reddy et al. 2012; Clements et al. 2023), but slightly over half belong to a monophyletic subfamily endemic to Madagascar (Vanginae). The Malagasy Vanginae have diversified into a spectacular array of forms, misleading early taxonomists who initially classified them as members of numerous other bird families (Yamagishi et al. 2001; Johansson et al. 2008; Reddy et al. 2012). Vangas are understudied relative to other avian radiations, notably the Galapagos finches and Hawaiian honeycreepers, and differ significantly from these clades in many aspects of natural history.

Vangas are all insectivores (some incorporating vertebrate prey), and have diversified primarily in terms of foraging strategy rather than diet: different species employ a diversity of maneuvers which can be broadly categorized as gleaning, sallying, and probing, with corresponding morphological specializations (Yamagishi and Eguchi 1996; Reddy and Schulenberg 2022). The Malagasy vangas are also a relatively old radiation, having colonized Madagascar over fifteen million years ago (Reddy et al. 2012; Oliveros et al. 2019). The diversity of biomes on this island continent has facilitated both ecological specialization and an exceptional degree of sympatry, with about fifteen species co-occurring in rainforest communities (Wilmé 1996). Prior studies which examined diversification of the Malagasy vangas found speciation rates largely consistent with the classic model of an early burst following ecological opportunity (Jønsson et al. 2012; Reddy et al. 2012). Using standard linear measurements of bill and body, (Jønsson et al. 2012) found that relative trait disparity also fit an early burst, but detected a marginal secondary burst in both relative disparity and speciation rate corresponding to the origin of derived “probing” foraging behaviors.

In this study we examine patterns of morphological diversification in the Vangidae, focusing on anatomical regions closely tied to foraging behavior: the bill, cranium, and feet. The bill is the ecomorphological trait most frequently studied in birds, being closely tied to diet and foraging behavior (Zusi 1993; Dehling et al. 2016; Pigot et al. 2016; Cooney et al. 2017; Krishnan 2023; Mosleh et al. 2023). The bill is continuous with the rest of the cranium, together forming the skull. The cranium itself includes features reflecting variation in musculature, vision, and neuroanatomy (van der Meij and Bout 2008; Navalón et al. 2020; Eliason et al. 2021). We applied geometric morphometric methods, following recent papers that have done so in birds (Cooney et al. 2017; Tokita et al. 2017; Felice et al. 2019; Eliason et al. 2020; Navalón et al. 2020; Vinciguerra et al. 2024), which use a series of homologous landmarks and semilandmark curves to describe complex variation in shape (Goswami et al. 2019). Avian pedal morphology is far less studied, although the diversity of hindlimb anatomy in birds has known relationships to locomotory mode (Miles and Ricklefs 1984; Abourachid et al. 2017; Falk et al. 2021; Dickinson et al. 2023). We used linear measurements of each pedal bone to describe variation in overall foot shape.

A notable feature of the Malagasy vanga radiation is the presence of a handful of taxa which have evolved especially divergent and sometimes unusual morphologies associated with specialized foraging strategies. Examples include the extremely long, decurved bill of *Falculea palliata*, specialized for probing in cavities in search of prey, the massive, deep, and strongly hooked bill of the sallying predator *Euryceros prevostii*, and modified foot proportions in the terrestrial *Mystacornis crossleyi* and nuthatch-like, tree-creeping *Hypositta corallirostris* (Johansson et al. 2008; Reddy and Schulenberg 2022). The presence of such extremes is a fundamental component of the radiation, but also has the potential to disproportionately drive signal or alter the results of analyses.

We used a range of geometric morphometric and phylogenetic comparative methods to explore the tempo and mode of morphological evolution in vangas and address several fundamental questions in the study of adaptive radiation. We took a comparative approach, contrasting the radiation of Malagasy vangas with the rest of the Vangidae, to evaluate the degree to which ecological opportunity in Madagascar has facilitated exceptional diversification. First, we assessed whether overall morphological diversity is in fact higher in the Malagasy clade than their mainland relatives. Second, we asked whether stronger integration or modularity is associated with morphological diversification across these clades. Third, we examined how rates of evolution have varied through time across Vangidae. Throughout, we evaluated how including multiple anatomical traits shifts our view of diversification patterns and assessed the evidence for correlated evolution. We also sought to evaluate the role of extremely divergent morphologies in driving our analyses of trait diversification.

## Methods

### Data collection

We obtained morphometric data from microCT scans of museum specimens, both alcohol-preserved (fluid) whole specimens and stuffed round skins. These specimen types are preferred because they preserve the rhamphotheca, the keratinous sheath which interacts directly with the environment (Cooney et al. 2017; Eliason et al. 2020; Chhaya et al. 2023). Fluid specimens are preferred because the entire body is preserved, but to improve our taxonomic sampling we included round skins, which contain only partial skeletons (usually the anterodorsal portion of the skull and distal portions of the limbs). Specimens are from the Field Museum of Natural History, American Museum of Natural History, and Yale Peabody Museum, including some scanned as part of the openVertebrate (oVert) project (Blackburn et al. 2024) and downloaded from Morphosource. Specimen and scan details can be found in Supplementary Table 1. In total, we obtained scans of 75 specimens, with most species represented by more than one specimen (Table S1). Our morphometric dataset included all genera and 34 of the 41 recognized species of Vangidae (Younger et al. 2019; Clements et al. 2023), though we were only able to obtain foot measurements for 30 species. Body mass was taken from the AVONET dataset (Tobias et al. 2022) except for *Schetba (rufa) occidentalis*, which was found in (Safford and Hawkins 2013).

We segmented each scan using 3D Slicer (Fedorov et al. 2012; Kikinis et al. 2014), then cleaned and smoothed all meshes in Autodesk Meshmixer (Autodesk Inc. 2018), and placed landmarks on each module in Stratovan Checkpoint (Stratovan Corporation 2022). Our landmark dataset consists of 214 landmarks, including 13 stationary landmarks and 9 curves (Figure 1, Table S2, see supplemental methods for further details). Our landmarking scheme follows that of other studies, particularly (Eliason et al. 2020), who also included the lower bill, and the landmark and sliding semi-landmark scheme of (Felice and Goswami 2018), with our choice of cranial landmarks constrained to those regions consistently preserved in round skins. We placed landmarks only on the left side to avoid unnecessarily increasing the dimensionality of the data (Cardini 2017). For the hindlimbs, we digitally measured in 3D Slicer the length of 12 bones from one foot of each specimen: the tarsometatarsus, first metatarsal, and all but the ultimate (ungual) phalanges of each digit (Figure 1, Table S2).

**Figure 1.**
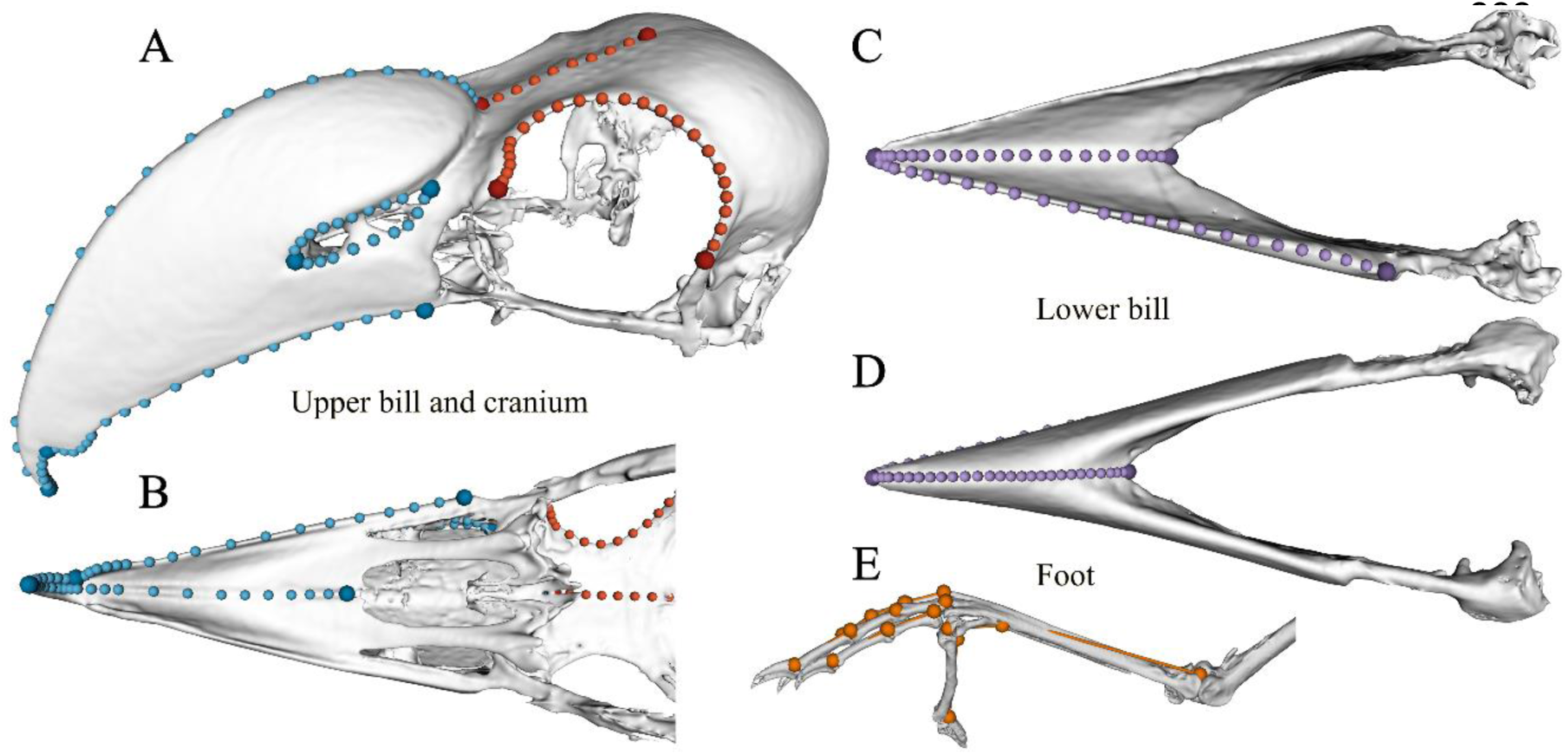
3-dimensional landmark configurations and linear measurements of pedal bones used in this study. Stationary landmarks are shown as larger, darker points, located mostly at the end of each semilandmark curve. Left: A) dorsolateral and B) ventral view of the upper bill (blue) and cranium (red). Right: C) dorsal and D) ventral view of the lower bill (purple), and E) linear measurements were taken between each pair of points on the pedal bones (orange).

All landmarks were aligned using a generalized Procrustes analysis (GPA) in the R package *geomorph* (Gower 1975; Rohlf and Slice 1990; Baken et al. 2021; Adams et al. 2024); semilandmark curves were slid to minimize bending energy (Bookstein 1997). Prior to performing the GPA, we mirrored the left-side only landmarks using the mirrorfill function in the R package *paleomorph*, then deleted the mirrored landmarks (Cardini 2017; Lucas and Goswami 2017). The lengths of hindlimb bones were transformed into log-shape variables by dividing by the geometric mean and then taking the natural log of that ratio, following e.g. (Slater 2022; Roberts-Hugghis et al. 2023). For each dataset we checked that specimens of the same species clustered together in a principal component analysis (PCA), then averaged coordinates by species for all subsequent analyses.

Our morphological dataset comprised of four anatomical modules which we used for all analyses: upper bill, lower bill, cranium, and feet (Figure 1). The three skull modules (bill and cranium) are functionally and structurally discrete, and are consistent with prior work that identified the major modules of the avian skull (Felice and Goswami 2018), although the lower bill was not included in that study. We performed a separate GPA on each module to avoid averaging variance across modules, which is of particular concern when some regions project far from the center of the object, as with bird bills (Cardini 2019; Cardini and Marco 2022).

We used a time-calibrated phylogeny of the Vangidae produced using reduced-representation genomic data targeting ultraconserved elements (UCEs; Reddy et al. in prep). In brief, this analysis included all described species of the family Vangidae except for 5 species outside of Madagascar (78 taxa total). Given the high variability of phylogenetically informative sites in UCE loci, which can exacerbate divergence dating analyses, we pruned our dataset to include only the 1000 most informative loci (following (Chen et al. 2021)) and then randomly subsampled 100 loci to create 10 subsets. For each subset, we ran an entropy estimation script to identify regions with differential rates of evolution (i.e. conserved cores vs. variable flanking regions) and used PartitionFinder 2 (Lanfear et al. 2017) to identify sets of loci with similar rates that can be combined in our model estimates. In BEAST 2.5 (Bouckaert et al. 2019), we conducted analyses of each subset for 100 million generations using a fixed topology from our maximum likelihood analyses and calculating branch lengths to estimate divergence times using the partition scheme and model settings from the PartitionFinder results. We used calibration estimates from Claramunt & Cracraft (Claramunt and Cracraft 2015) which used the largest set of verified fossils to date, to fix the MRCA of the root (Vangidae + Platysteridae) as 27.36 Ma (95%HPD 22.71 -32.09). We compared our resulting divergence times to those of other recent analyses of passerines (Oliveros et al. 2019) and found them to be comparable and within the 95% confidence intervals.

### Patterns of morphological diversity

To visualize the major axes of shape variation in vangas we performed PCAs on the GPA-aligned landmarks and log-shape variables. We focused on standard, rather than phylogenetic, PCA, as this results in strictly orthogonal axes which are required as input for several downstream analyses (Revell 2009; Polly et al. 2013). We performed separate PCAs on each module and on the combined landmark dataset to assess how overall patterns of variation differed between anatomical regions.

Initial exploratory analyses identified several Malagasy taxa as extreme outliers in each module (see Results; Figure 1, Table 1). To visualize the effect of these outliers on the primary axes of variation, we performed additional PCAs in which we initially excluded the outlier taxa. The outliers were then projected into the ordination by multiplying their GPA-aligned landmark configurations or mean-centered log-shape variables by the eigenvectors from the initial PCA; this is referred to as a posthoc rotation. Most subsequent analyses were performed on both the full set of Malagasy vangas and with either one or two outliers excluded to assess the degree to which our results were driven by the presence of these extreme morphologies.

**Table 1.**
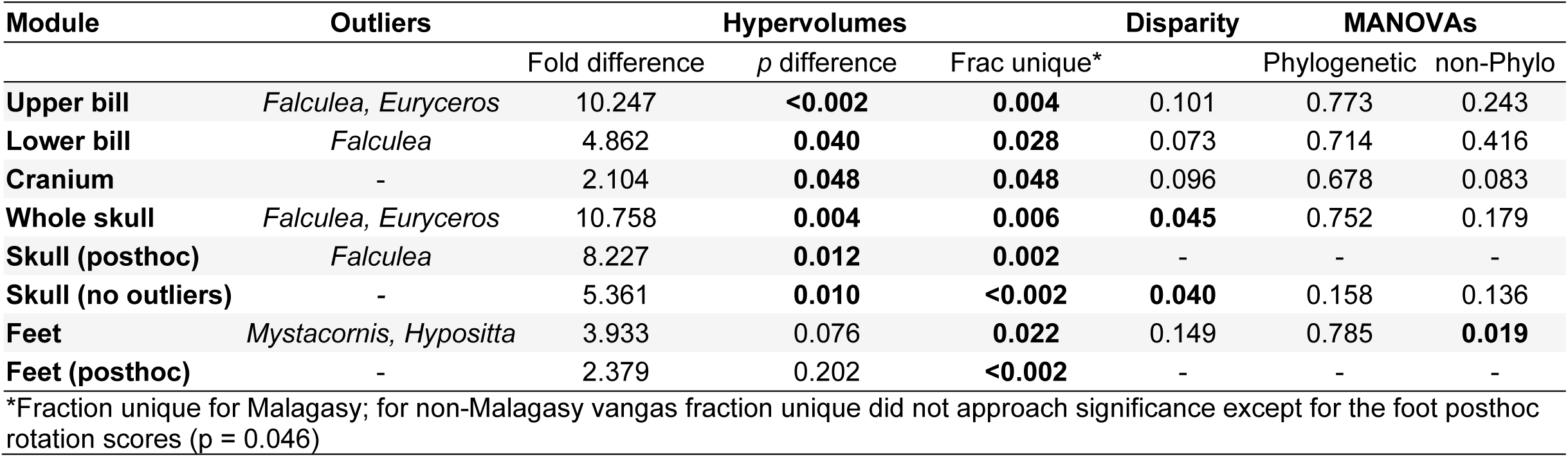
Morphological diversity of the Malagasy vs. non-Malagasy vangas by anatomical region. Quantified using kernel density hypervolumes from the first 3 PC axes for each module, and disparity and MANOVAs from the full landmark or linear measurement dataset. Outliers are identified from PC axes. Fold difference is the magnitude of difference in hypervolume size; fraction unique is percentage of hypervolume unique to the subclade. Significance of difference between clades (p value) is reported for each statistic.

To quantify differences in morphological diversity between the Malagasy versus non-Malagasy vangas and between foraging categories, we used kernel density hypervolumes, overall disparity (based on Procrustes variance), and multivariate analysis of variance (MANOVA). Foraging categories follow (Reddy et al. 2012). Kernel density hypervolumes have the benefit of accounting for the presence of holes in the morphospace, such as those created by outliers, but are limited by the hypervolume space becoming sparse as the dimensionality of the data increases (Blonder 2016). On the other hand, overall disparity does not account for holes in morphospace but has the benefit of using the full dimensionality of the dataset. We used the hypervolume_gaussian function in the R package *hypervolume* (Blonder 2016; Blonder et al. 2018) to calculate hypervolumes for the first three PC axes for each module as well as the entire skull dataset (Figure 3). We set the quantile boundary to 0.8 for skull modules and 0.65 for the feet, which was the most conservative threshold at which the outlier taxa remained consistently included. To evaluate whether differences in hypervolume size between the Malagasy and non-Malagasy clades were significant, and whether either clade occupied unique regions of morphospace, we performed a permutation test of group membership using the hypervolume_permute function and computed overlap statistics.

We compared the degree of overall disparity using the morphol.disparity function in *geomorph* (Zelditch et al. 2012). We also used a multivariate analysis of variance (MANOVA) to determine whether there were overall differences in shape, implemented using the functions procD.lm and procD.pgls (for phylogenetic analyses of variance) in *geomorph* (Adams 2014a; Adams and Collyer 2016). We additionally performed individual ANOVAs for each of the twelve foot bones.

We estimated the degree of phylogenetic signal for each module and compared its strength both between clades and between modules, using the physignal.z function in *geomorph*. This method finds the lambda transform which maximizes likelihood using residual randomization in a permutation procedure (RRPP; (Collyer and Adams 2018, 2024; Collyer et al. 2022)). Estimating phylogenetic signal for high-dimensional data is not a fully resolved challenge; current best practice suggests using phylogenetically aligned PCA (PACA) to reduce the number of data dimensions below the number of observations and generate a smaller number of variables which maximize the phylogenetic signal present in the data (Collyer et al. 2022). While the method of using axes which maximize phylogenetic signal will naturally tend to bias results towards finding significant signal in the data, our questions are focused on whether there are differences between clades. We selected the number of PACA dimensions to include for each data subset based on the flattening of the partial RV coefficient, using a cutoff of 99.5% of covariation explained between data and phylogeny. This was implemented for each data subset by calculating the number of dimensions to include from the PACA, setting the PAC.no argument equal to this value, and selecting the “front” lambda optimization method, with ten thousand iterations used for significance testing. Including only a subset of axes even after dimensionality reduction was necessary because axes explaining essentially zero covariation introduced substantial noise into the analysis which heavily influenced the results.

### Integration and Modularity

The degree of integration can be quantified both within and between anatomical modules, and these quantities together relate to modularity, often measured as the ratio within to between module integration. For within-module integration we calculated the relative eigenvalue index using the integration.Vrel function in *geomorph* and accounting for phylogeny (Pavlicev et al. 2009; Conaway and Adams 2022). This index measures integration as relative eigenvalue variance, as variables which covary more can be concentrated onto fewer eigenvalues, standardized by the number of traits. We compared the strength of integration between modules and clades with the function compare.ZVrel, which performs two-sample z tests of effect size (Conaway and Adams 2022). We compared the strength of integration both for the skull modules alone, and between all modules, with those taxa not represented in the foot dataset removed.

Integration of different modules was quantified using a phylogenetic partial least squares analysis (PLS) implemented using the phylo.integration function in *geomorph* (Bookstein et al. 2003; Adams and Collyer 2016). We conducted PLS analyses for the adjacent skull modules (upper bill and skull, upper and lower bill) as well as between each skull module and feet. To determine whether there are significant differences in the strength of integration between the two clades, we used the function compare.pls to test for differences in effect sizes between the Malagasy and non-Malagasy vangas.

To compare the degree of modularity between the Malagasy and non-Malagasy vangas we calculated the covariance ratio (CR) coefficient for each pair of modules using the phylo.modularity function in *geomorph* (Adams 2016; Adams and Collyer 2019). This calculates the ratio of within-versus between module covariance, which is compared to a null CR distribution based on permuting landmark group membership to test for significance, while accounting for covariation due to phylogenetic structure. It also calculates Z scores that are used to test for a difference in the strength of modularity between the two clades.

### Evolutionary Rate Analyses

To assess how rates of morphological evolution have changed through time we compared the fit of a series of local tree transforms using five different scaling parameters, as implemented in BayesTraits V4.0.1 (Venditti et al. 2011; Pagel et al. 2022). The transforms are termed kappa, lambda, delta, node and branch. Kappa, lambda, and delta transforms have a default value of 1, indicating that the rate of evolution is generally constant through time as the true branch lengths accurately represent patterns of trait evolution; deviations represent different trends of rate change through time (Pagel 1999) A kappa > 1 stretches longer branches whereas a kappa < 1 stretches shorter ones, such that smaller values of kappa are consistent with a more punctuated mode of evolution. Lambda is a measure of the degree of phylogenetic signal in the data, with lambda < 1 indicating increasingly weak correspondence between trait values and evolutionary relatedness. A delta < 1 increases the length of branches towards the root of the phylogeny relative to the tips, consistent with an early burst of evolution, whereas delta > 1 increases branch lengths towards the tips, consistent with accelerating rates of evolution through time. The node transform scales all descendant branches of that node by some value, effectively increasing or decreasing the evolutionary rate σ^2^ for that portion of the phylogeny, while a branch transform applies to a single branch and indicates a change in the mean value of a trait for the descendant clade.

For each transform we tested the fit of three rate shift placements – 1) at the root of the Malagasy clade (equivalent to early burst); 2) at the root of the “derived” clade of foraging behaviors; 3) multiple shifts occurring elsewhere on the phylogeny. For the first two, we specified the node at which the transform would occur using the *LocalTransform* command. For the third, we used a reversible-jump Markov chain Monte Carlo (rjMCMC) approach (*RJLocalTransform* option in Bayestraits) to determine for each transform the location and number of shifts that best fit the data. Finally, we implemented the “Fabric” model (Pagel et al. 2022), which uses rjMCMC to simultaneously detect directional (branch transform) and evolutionary rate changes (node transform). See Table S9 for a complete list of all tested models. Because kappa, lambda, and delta transforms stretch some branches relative to others, they cannot be applied to tips. By default, BayesTraits restricts these transforms to nodes of 10 taxa or more; due to the relatively small size of our phylogeny we changed this restriction to nodes of 5 taxa or more, so as to not force changes in evolutionary mode towards the root of the tree. Branch and node transforms, however, were left free to occur at any point on the tree.

We used these models to evaluate the tempo and mode of evolution in a total of five trait datasets: whole skull, whole skull with post-hoc rotation, bill (upper + lower), feet, and body mass. For all multivariate trait datasets we used the principal component axes that summarized 95% of the total shape variation to reduce data dimensionality. This resulted in thirteen PC axes for the skull dataset, fourteen for the skull with posthoc rotation, nine for the bill, and seven for the feet.

For each model we ran five independent Markov chains each for at least eleven million generations, sampling every thousand generations and discarding the first one million generations as burn-in. We used Tracer v1.7.2 to check for stationarity and convergence (Rambaut et al. 2018). Some reversible-jump models failed to converge after eleven million generations; these we ran for fifty or one hundred million generations after burn-in, reducing the sampling frequency to retain a constant number of samples. The fit of each model was compared to a null model of constant rates using Bayes factors from a stepping-stone sampler with one thousand steps each run for fifty thousand generations. In addition to calculating Bayes factors, we identified the location, magnitude and support for rate shifts, and visualized branch-specific rates across the phylogeny for the best-supported models, using the R packages *BTprocessR* (Ferguson-Gow 2020) and *ggtree* (Yu et al. 2018).

As a complementary means of understanding the rate of trait diversification through time, we performed a disparity-through-time analysis on each trait dataset using the function dtt in the R package *geiger* (Harmon et al. 2003; Slater et al. 2010; Pennell et al. 2014). In this analysis relative subclade disparity (based on pairwise Euclidean distances between species) is calculated as the ratio of subclade to total clade disparity. Mean relative disparity over time is then calculated as the average among all subclades present at each point, and this observed relative disparity is compared to a null BM model of trait diversification. Smaller values of relative subclade disparity indicate a greater partitioning of shape variation among clades, consistent with an early burst of morphological evolution, whereas high values indicate that subclades encompass a relatively large portion of the total clade’s morphospace.

We also compared the net rate of morphological evolution for the entire landmark dataset, for each module, and for individual pedal bones between the Malagasy and non-Malagasy vangas and between foraging modes, using the compare.evol.rates function in *geomorph* (Adams 2014b; Denton and Adams 2015). This analysis finds the net rate of evolution under a Brownian motion model and calculates the ratio of rates for two or more groups. Both permutation-and simulation-based methods are available for assessing the significance of the rate ratio; we evaluated the results of both options (Adams and Collyer 2018).

## Results

### Patterns of morphological diversity

Our analyses confirmed that Malagasy vangas are much more morphologically diverse in the measured traits than non-Malagasy vangas. PCAs (Figure 2) show that across all four modules, the Malagasy vangas occupy a far greater total spread of morphospace along the primary axes of variance. As expected, some species with extreme morphologies appear to be outliers in how distant they are from other species and drive much of the variation seen on PC1 and PC2, although numerous Malagasy vangas fall well outside the morphospace of the non-Malagasy vangas (Figure 2). *Falculea* (Sickle-billed Vanga), is highly divergent in both upper and lower bill shape (Figure 2c-d), while *Euryceros* (Helmet Vanga), is an outlier in upper bill only Figure 2c). In the feet, *Mystacornis* (Crossley’s Vanga) and *Hypositta* (Nuthatch-Vanga) were outliers, with the difference between them defining the primary axis of variation (Figure 2f). *Mystacornis* has a proportionally elongated tarsometatarsus and shortened hallux, whereas the reverse is true in *Hypositta*. Although the Malagasy vangas were more diverse than the non-Malagasy vangas in cranial shape, no taxa were particular outliers Figure 2e). The Malagasy vangas also occupied a greater region of morphospace on subsequent PC axes (Figure S1).

**Figure 2.**
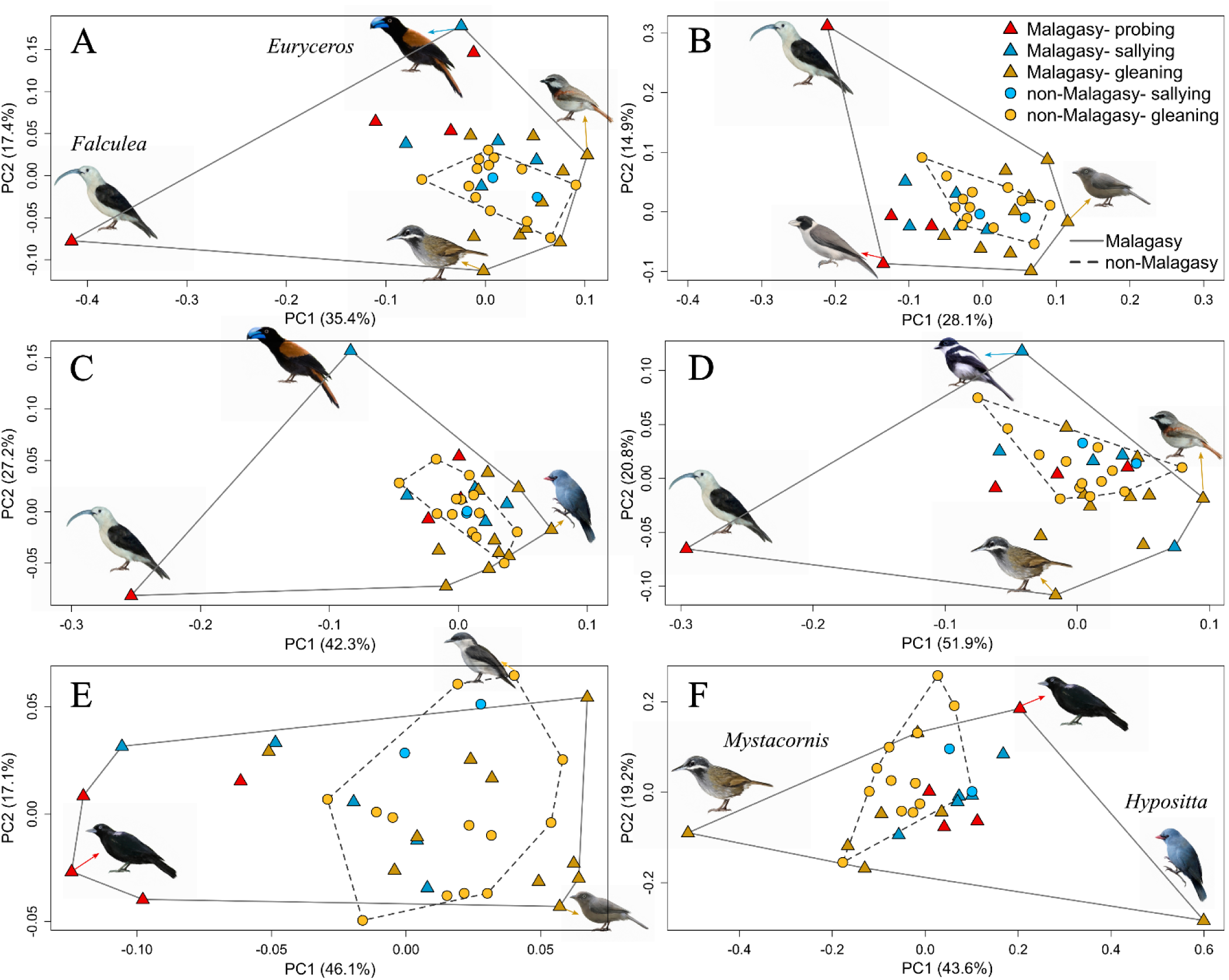
Morphological diversity in vangas. Principal components 1 and 2 from PCAs of GPA-aligned landmark configurations for the skull and log-shape variables for the feet. Top row: A) whole skull and B) whole skull posthoc rotation scores, showing differences in the primary axes of variance when outliers are excluded. Middle: C) upper bill and D) lower bill. Bottom: E) cranium and F) feet. Triangles represent Malagasy vangas, while circles represent non-Malagasy vangas. Points are colored by foraging category, with red for probing, blue for sallying, and yellow for gleaning; darker shades are used for the Malagasy vangas. Convex hulls are drawn around each clade, solid (Malagasy) and dashed (non-Malagasy). Inset birds represent the extremes of shape; outliers are labeled.

We performed PCAs with posthoc rotation of these outliers on the whole skull and foot datasets. In both, the results show that the outliers did substantially alter the loadings of variables on the primary axes of variance, changing the general patterns that emerged for the clades as a whole (Figure 2b, Figures S1 and S2). To evaluate the role these outliers play in driving the overall diversity of the Malagasy radiation, we compared for each test the Malagasy vangas as a whole, the Malagasy vangas with outliers removed, the Malagasy vangas using posthoc rotation scores (where relevant), and the non-Malagasy vangas.

Malagasy and non-Malagasy vangas differed in morphological diversity, mean shape, or both for all measured traits (Table 1). Our analyses of morphospace hypervolumes found that the Malagasy vangas occupied significantly more morphospace than the non-Malagasy vangas for all skull modules but not the feet, and that a significant fraction of the morphospace was unique to the Malagasy vangas for all trait datasets (Table 1, Figures 3 and S3). Overall disparity was always greater in the Malagasy vangas, but differences were only significant for the whole skull (Table 1). MANOVAs indicated that average shape was significantly different between the clades only for the feet (Table 1). ANOVAs of each pedal bone indicated that these differences are spread across the first three digits (Table S3).

**Figure 3.**
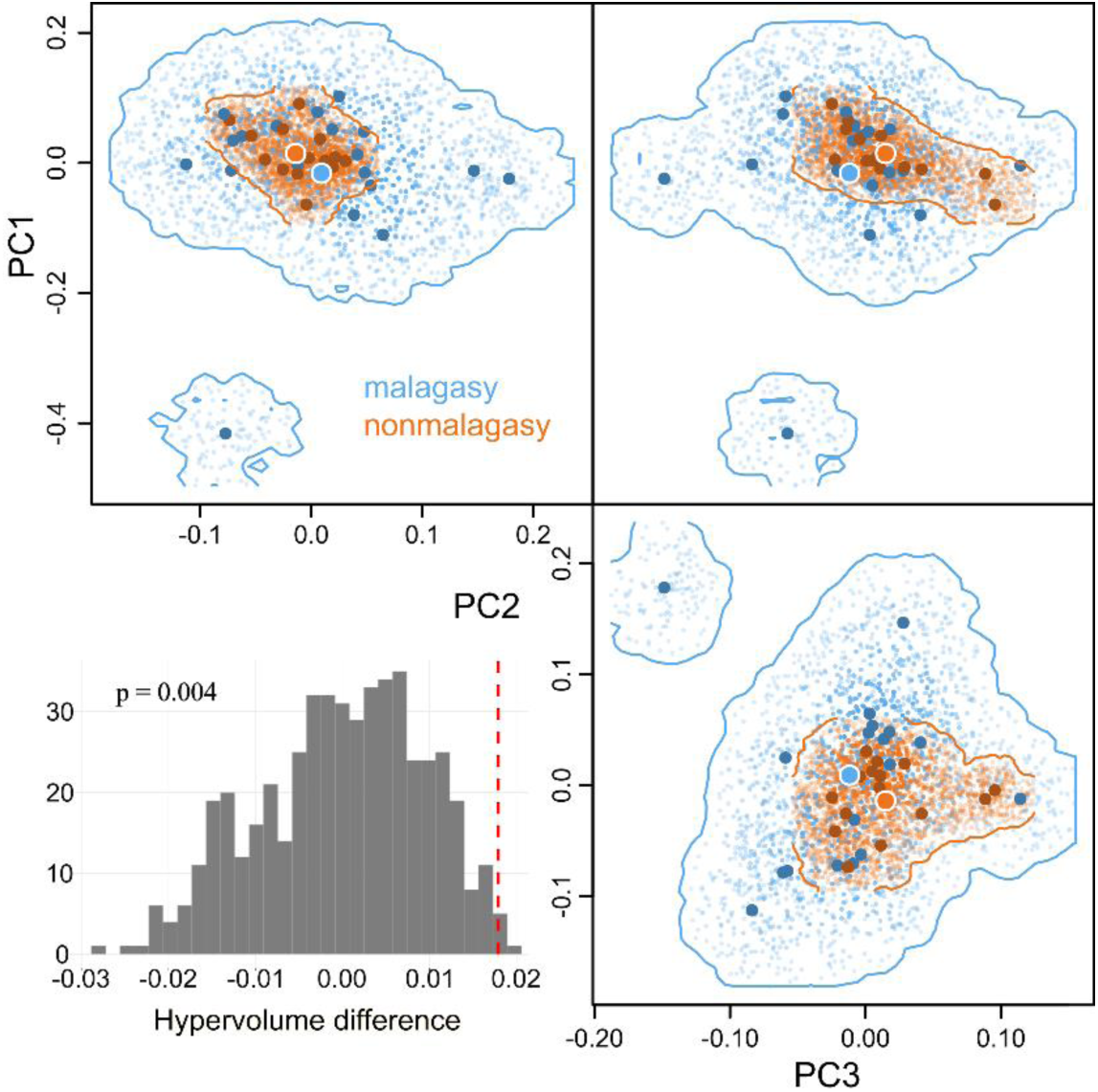
Morphospace hypervolumes of the skull for Malagasy (blue) and non-Malagasy (orange) vangas. Hypervolumes are for the first three PC axes from the whole skull landmark dataset. Large, dark points represent the observed point for each species, and the small points the density distribution of the hypervolume. The two largest points with white borders indicate the centroid of each clade. The inset histogram is the null distribution of hypervolume differences from the permutation test, with a dashed line indicating the observed difference in volume

Vangas in the three foraging categories occupy distinct regions of morphospace and differ in their degree of overall morphological disparity. In the whole skull and cranium, gleaners and probers do not overlap on the first two PC axes, with salliers occupying an intermediate position (Figure 2). MANOVAs indicate that these differences in overall shape are significant for all skull modules, but only when not accounting for phylogenetic signal (Table S4). Probers consistently had greater disparity than the other foraging classes (Table S4). For foot shape, gleaners dominate the primary PC axes because both outliers are in this category (Figure 2). Foraging classes were also significantly different in mean foot shape, though not in disparity, but only when analyzed using posthoc rotation scores (Figure S2, Table S4).

We detected significant phylogenetic signal for most modules and subclades, and differences between categories were nearly all statistically significant (Table S5). In the Malagasy vangas, phylogenetic signal was highest in the cranium and lowest in the bill: the upper bill in Malagasy vangas was the only module with no phylogenetic signal. In contrast, phylogenetic signal in the non-Malagasy vangas was highest in the upper bill. For the lower bill, phylogenetic signal was notably highest when only *Falculea* was removed from the dataset. Phylogenetic signal in foot shape was lowest in the Malagasy vangas with outliers removed, and highest in the non-Malagasy vangas.

### Integration and modularity

Our outliers (*Euryceros* and *Falculea*) were not equally divergent across anatomical regions. Because this might impact clade-wide patterns of trait covariation, we explored the impact of removing them from the Malagasy vanga dataset one at a time, as well as together in our analyses of integration and modularity.

Within-block integration (eigenvalue dispersion) was consistently but not significantly higher in the Malagasy vangas across modules (Table S6). Removing both *Falculea* and *Euryceros* from the Malagasy dataset always decreased integration in the remaining Malagasy vangas, usually below that of the non-Malagasy clade. However, removing only *Euryceros* from the whole skull and upper bill datasets left the remaining Malagasy vangas significantly more integrated than the non-Malagasy, as well as than the Malagasy with both outliers removed (Table S6).

The upper bill and cranium were more integrated in the Malagasy vangas, while the upper and lower bill were more integrated in the non-Malagasy vangas, but in both comparisons removing both outliers reversed this pattern (Table S7). Plotting the scores from the first axes of covariance revealed that both *Euryceros* and *Falculea* deviate substantially from the dominant axes of covariance for the rest of the clade (Figure 4). As with eigenvalue dispersion, few differences were statistically significant. For the upper bill and cranium, removing only *Falculea* increased integration; the remaining Malagasy vangas are significantly more integrated than when both outliers are removed (p=0.04). For the upper and lower bill, Vangidae as a whole show significantly less integration than the Malagasy vangas with both outliers removed, but significantly more than the non-Malagasy vangas or the Malagasy without *Falculea* (Table S7). Nearly all tests for integration between the feet and skull modules found no significant covariation with the exception of the cranium-plus-upper bill, which were significantly integrated with foot shape only in the non-Malagasy vangas (p=0.02).

**Figure 4.**
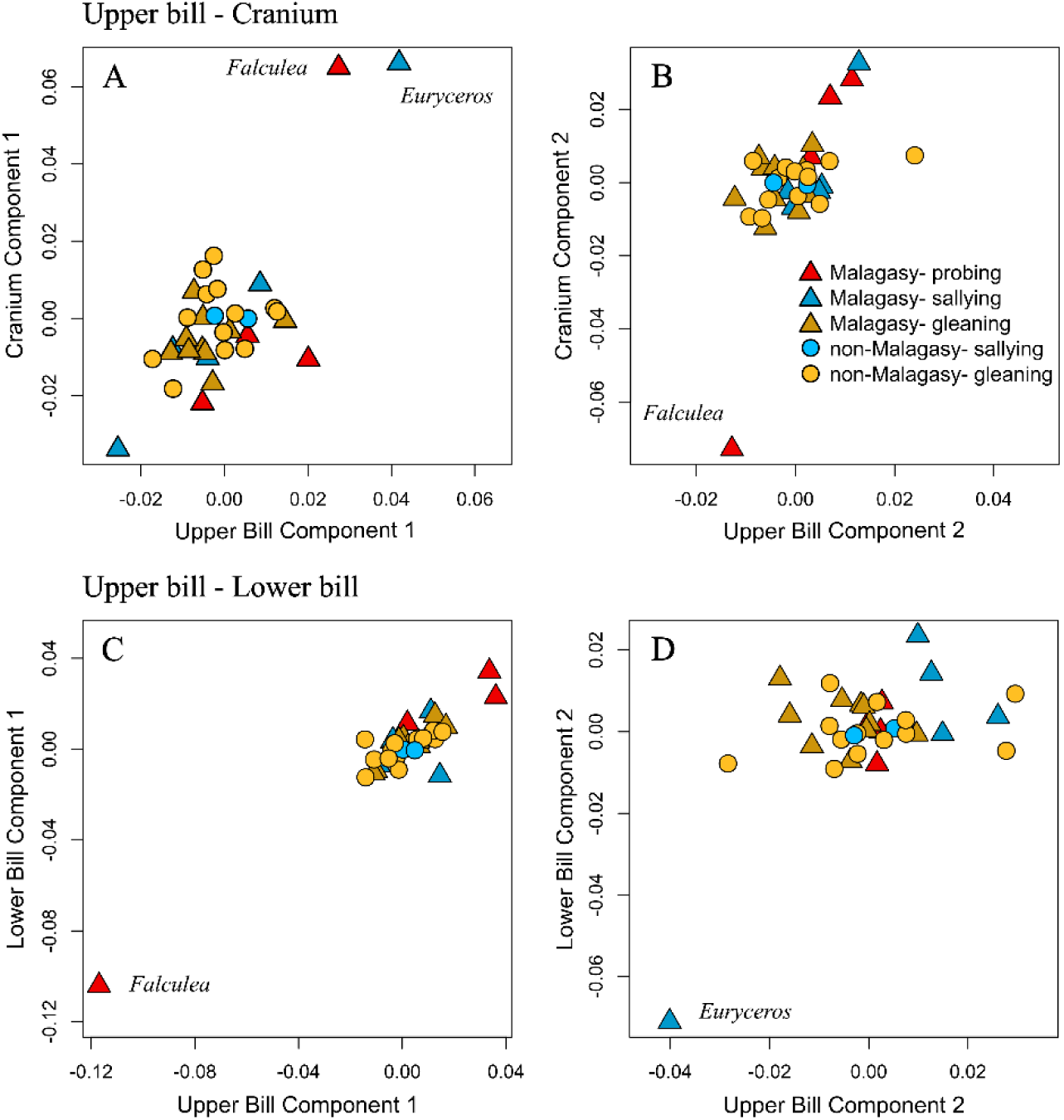
Skull shape integration in vangas. Scores from the first two pairs of PLS axes testing integration of the upper bill and cranium (top) and upper and lower bill (bottom). These are similar to PCA plots, but show the primary axes of mutually predictive variance for each pair of anatomical modules. PLS component one (A, C) and two (B, D). Triangles represent Malagasy vangas, while circles represent non-Malagasy vangas. The points are colored by foraging category, with red for probing, blue for sallying, and yellow for gleaning; darker shades are used for the Malagasy vangas. The positions of anatomical outliers (*Euryceros*, blue and *Falculea*, red) are indicated on each plot.

Modularity was consistently higher in the Malagasy than the non-Malagasy vangas; this result was highly significant for the upper and lower bill (p<0.001) but not for the upper bill and cranium (Table 2). In both cases, removing *Falculea*, *Euryceros*, or both always decreased the degree of modularity in the remaining Malagasy dataset. Removing *Euryceros* or both outliers resulted in the remaining Malagasy vangas showing significantly lower modularity than the non-Malagasy vangas, while removing *Falculea* alone had a more variable impact. Examining the distribution of null covariance ratios from the resampling procedure for each test suggested that wide variation in the mean and standard error of these distributions between datasets may explain at least some of the surprising variation in significance test results (Figure S4).

**Table 2.** Pairwise comparisons of the strength of skull modularity between Malagasy and non-Malagasy vangas. Modularity is measured by the covariance ratio (CR). Z_CR_ is the strength of modularity for each clade, with a more negative Z score indicating greater modularity – the Malagasy vangas (with outliers included) are the most modular. Pairwise comparisons of modularity between clades (Malagasy, non-Malagasy, and Malagasy with outliers removed), with Z scores above and p values below the diagonal. Significant differences are bolded.

| Upper Bill vs Cranium | Malagasy | non-Malagasy | Malagasy (no outliers) | Malagasy (no Falcula) | Malagasy (no Euryceros) |
| --- | --- | --- | --- | --- | --- |
| Z<sub>CR</sub> (Modularity) | -15.523 | -7.946 | -6.047 | -9.812 | -5.280 |
| Malagasy | - | 0.057 | **3.641** | **4.955** | **12.567** |
| non-Malagasy | 0.954 | - | **2.696** | **3.139** | **6.778** |
| Malagasy (no outliers) | **<0.001** | **0.007** | - | 0.084 | **4.521** |
| Malagasy (no Falcula) | **<0.001** | **0.002** | 0.933 | - | **6.932** |
| Malagasy (no Euryceros) | **<0.001** | **<0.001** | **<0.001** | **<0.001** | - |
| Upper Bill vs Lower Bill |  |  |  |  |  |
| Z<sub>CR</sub> (Modularity) | -16.221 | -14.223 | -10.718 | -12.129 | -14.109 |
| Malagasy | - | **7.600** | **4.127** | **6.466** | **11.696** |
| non-Malagasy | **<0.001** | - | **2.746** | 0.208 | **12.625** |
| Malagasy (no outliers) | **<0.001** | **0.006** | - | **2.336** | **9.090** |
| Malagasy (no Falcula) | **<0.001** | 0.835 | **0.019** | - | **10.751** |
| Malagasy (no Euryceros) | **<0.001** | **<0.001** | **<0.001** | **<0.001** | - |

Overall, we did not find a consistent relationship between the morphological diversity of the Malagasy vangas and either integration or modularity; instead we found that the results were broadly sensitive to the presence of anatomical outliers. Removing *Falculea* decreased within-module and bill integration, but increased integration between the bill and skull; it also slight decreased modularity. In contrast removing *Euryceros* increased within-module integration of the upper bill and integration of the upper and lower bill, but decreased integration between the bill and skull, and strongly decreased modularity.

### Rates of trait evolution

For all traits except body size, our BayesTraits analyses found very strong evidence in support of more complex, rate-variable models over a constant rate of morphological evolution (Table S9). The results for all three skull/bill datasets were essentially the same: we focus here on the full skull without posthoc rotation of outliers but results from all three can be found in the Supplemental material. The best-supported model for rates of vanga skull evolution was a reversible-jump branch transform, where a scalar was applied to certain tips, indicating a strong directional trend in shape in those taxa. Five branch scalars were found in greater than 85% of trees from the posterior, with consistent probabilities and magnitudes across five runs (Figure 5, Table S10). Shifts occurred in *Falculea* and *Euryceros* 100% of the time, with ∼20-25-fold increases, while smaller shifts usually occurred in *Xenopirostris* and two members of the African genus *Prionops* (Figure 5, Tables S10-11). The best-supported model for foot shape was a reversible-jump delta transform (Table S9). This model found a delta transform of 1.76 at the root of all Vangidae in 100% of posterior trees, indicating a family-wide trend of accelerating foot evolution through time (Table S10). An additional two delta transforms were detected in slightly under half of sampled trees: a second increase in delta of 1.18 at the ancestor of *Hypositta* plus the remaining Malagasy vangas, followed by a decrease in delta of 0.90 at the subsequent node (excluding *Hypositta*) (Figure S5). For body mass, we found moderate support for a node transform at the origin of the “derived” clade (Bayes factor = 2.78), indicating an increase in rates of size evolution in this group (Table S9).

**Figure 5.**
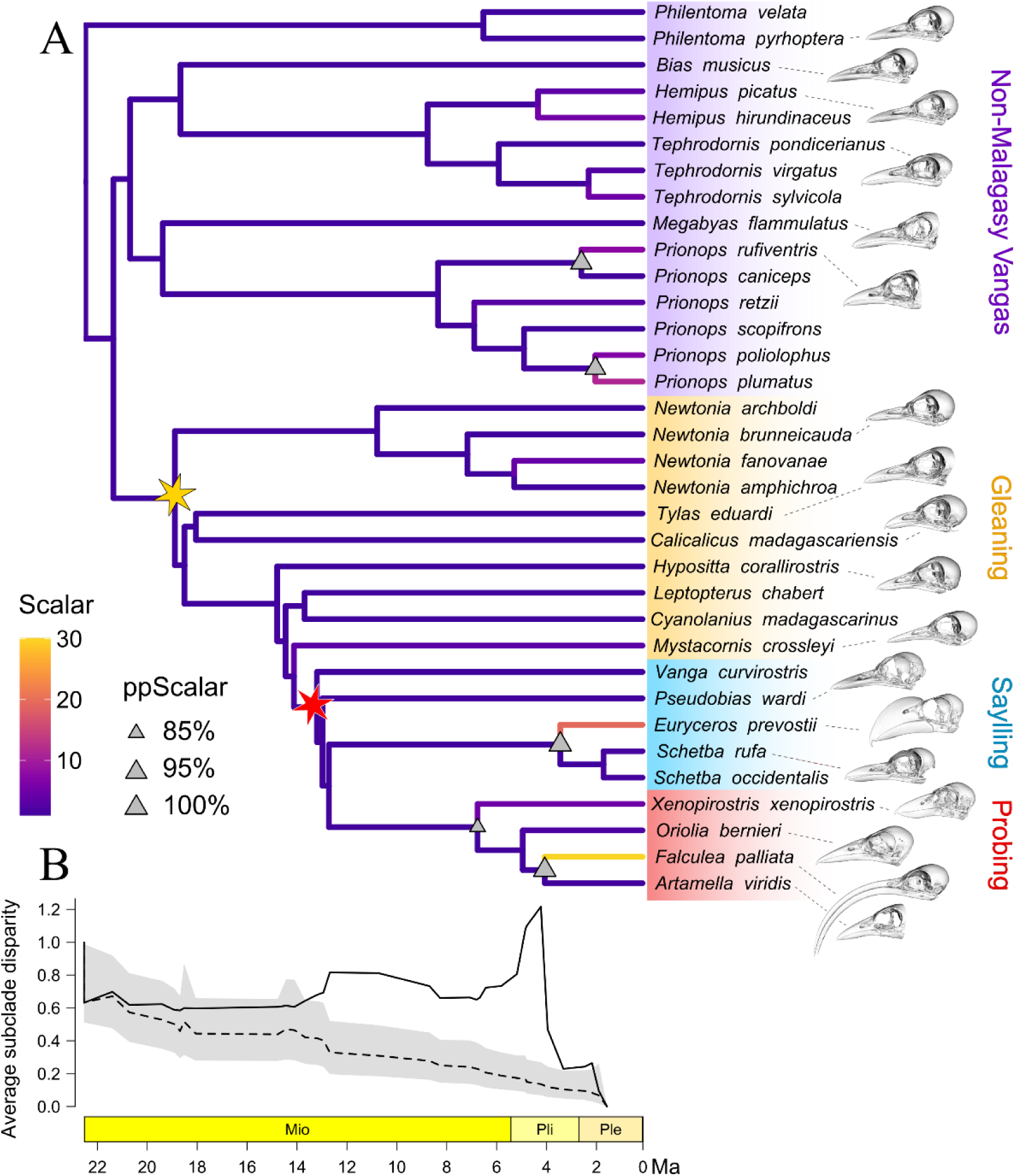
Rates of skull shape evolution in vangas. A) Phylogeny of Vangidae with branches colored by magnitude of directional shifts. The best-supported model suggests that large directional shifts in individual taxa have collectively generated the exceptional degree of morphological diversity observed in the Malagasy vangas. Triangles indicate shifts detected in over 85% of posterior trees, and stars indicate the colonization of Madagascar (yellow) and origin of the “derived” subclade (red). B) Disparity through time plot, with solid line indicating observed disparity, dashed line simulated disparity under constant rates, and the shaded area the 95% confidence interval. Average subclade disparity was elevated through the diversification of the derived clade, peaking just before *Falculea* split from its sister species *Artamella*.

Our dtt results are largely congruent with our BayesTraits analyses, showing that average subclade disparity in skull shape does not conform to Brownian motion expectations in the Malagasy vangas (Figure 5). Subclade disparity stayed within BM expectations until a sharp increase about 12 mya, at the common ancestor of *Euryceros* and the probing clade. It then stayed high before spiking about 4 mya, just before *Falculea* and *Euryceros* split from their respective sister taxa, then declined sharply towards the present. Foot shape, in contrast, showed a steady decrease in disparity through time consistent with the null expectation of a Brownian motion model of morphological evolution (Figure S6).

The net rate of multivariate trait evolution was 2-4x higher in the Malagasy vangas for each module except the cranium and for the whole skull, but removing the two outliers eliminated this difference (Table 3). The net evolutionary rate also differed significantly between vangas using different foraging behaviors (Table 3). The small but morphologically disparate probing clade consistently had the highest rates of skull evolution, up to 10.7x that of gleaners in the upper bill. However, the statistical significance of these differences varied by method. Rate differences were highly significant when tested using phylogenetic simulation (Denton and Adams 2015), but not when using the permutation test (Table S12) (Adams and Collyer 2018). The difference between the two methods might be a consequence of permutation being more sensitive to the presence of outliers.

**Table 3.** Differences in the net rate of evolution for each anatomical region between clades (Malagasy versus non-Malagasy vangas) and between foraging categories (probing, sallying, and gleaning). Values are rate difference (fold) and p values from the simulation procedure. Significant differences are bolded.

| Module | Malagasy vs. non-Mal | Probe vs. Glean | Probe vs. Sally | Glean vs. Sally |
| --- | --- | --- | --- | --- |
| Whole skull | **2.306 (0.001)** | **6.711 (0.001)** | **3.444 (0.001)** | **1.949 (0.001)** |
| Skull (outliers excluded) | 1.186 (0.160) | 1.424 (0.0829) | **1.881 (0.013)** | 1.321 (0.079) |
| Upper bill | **4.034 (0.001)** | **10.893 (0.001)** | **2.953 (0.002)** | **3.689 (0.001)** |
| Lower bill | **2.760 (0.001)** | **8.759 (0.001)** | **6.349 (0.001)** | 1.380 (0.250) |
| Cranium | 1.135 (0.462) | 1.537 (0.108) | 1.158 (0.618) | 1.328 (0.166) |
| Feet | **2.008 (0.003)** | 1.037 (0.919) | **2.375 (0.024)** | **2.290 (0.002)** |
| Feet (outliers excluded) | 1.112 (0.630) | 1.824 (0.056) | **2.284 (0.021)** | 1.252(0.373) |

## Discussion

### Malagasy vangas show exceptional ecomorphological diversity

Our analyses of morphological diversity in Vangidae confirmed that the Malagasy vangas are more morphologically diverse across all measured traits. This finding is consistent with a classic model of adaptive radiation in response to ecological opportunity, where the Malagasy vangas have diversified in traits which allow them to exploit a range of niches usually filled by other clades on the mainland. Some of the Malagasy diversification may have been facilitated by the evolution of new modes of foraging. Much of the novel bill shape diversity in the Malagasy vangas is found in the subclade in which first sallying, and then probing types of foraging maneuvers evolved. Except for two non-Malagasy taxa known to sally (both morphologically similar to the Malagasy *Pseudobias*), this behavior and probing are exclusive to the Malagasy clade, and in both a diversity of morphologies for different specialized types of sallying and probing have evolved. However, these foraging categories do not directly account for the foraging substrate or terrain used by different species, which likely drive locomotor adaptations and therefore pedal shape diversity. The most unique pedal morphologies are in gleaners with different locomotory demands: the terrestrial *Mystacornis*, and *Hypositta*, a vertical climbing gleaner of trunks and branches.

### Integration and modularity vary within the Malagasy vangas

The results of our analyses of integration and modularity were complex, suggesting that the interaction between integration, modularity, and evolvability in vangas is not straightforward. When we excluded anatomical outliers, the Malagasy vangas generally had statistically indistinguishable levels of integration and significantly lower modularity than the non-Malagasy vangas, suggesting some role for reduced independence at least between skull modules in their diversification. However, the presence of a few species with highly divergent morphologies among the Malagasy vangas had a profound effect on all our integration and modularity results. Including both *Euryceros* and *Falculea* qualitatively increased within-module integration, significantly increased integration of the upper bill and skull, and significantly increased modularity of the upper and lower bill over that of the non-Malagasy vangas.

The impact of including only one of the two bill shape outliers was highly variable, and this is likely due to their very different morphologies. Although *Falculea* has by far the most unique bill shape among all vangas, standing out in every one of our analyses, its overall patterns of trait covariation appear to be partially in line with the rest of Vangidae. This was evidenced by the results of the PCA in which we project the outlier taxa post-hoc into the transformed space defined by the patterns of covariation in the rest of Vangidae-*Falculea* remained an outlier in this space, albeit more on PC2 than PC1. Our PCAs also showed how the long, decurved bill of *Falculea* is similarly extreme in both upper and lower bill shape. In contrast, *Euryceros* has a distinctive upper bill – also highly decurved but domed and massive – but was not outlier in lower bill shape. This seems to lead to a unique pattern of trait covariation, resulting in *Euryceros* not appearing as an outlier on the primary axes of shape variation in the rest of Vangidae. We saw a similar result for both foot shape outliers, with *Mystacornis* and *Hypositta* not appearing as outliers in posthoc rotation plots. Our PLS plots show that for upper bill-skull integration, *Falculea* and *Euryceros* diverge from the rest of Vangidae in similar ways on the first pair of PLS axes, while for upper-lower bill integration, each is an outlier only on different axes, suggesting contrasting modes of difference. Overall, these results suggest a flexible relationship between integration, modularity, and diversification in vangas. The extremely decurved bill of *Falculea* requires maintaining tight integration between the upper and lower bill, whereas in *Euryceros* they are far more decoupled. A previous study that compared patterns of integration between the upper bill and cranium in two other adaptive radiations of birds, the Galapagos finches and Hawaiian honeycreepers, found higher levels of integration in these clades than in other passerine birds, but no general relationship between integration and rates of evolution (Navalón et al. 2020). They suggested that this relationship is likely to break down in older clades due to variation in selective pressures acting over many millions of years; our findings in Vangidae may reflect this. However, (Navalón et al. 2020) did not include the lower bill in their study. It would be interesting to compare our results with analyses including the lower bill in other birds such as the Hawaiian honeycreepers, which include several taxa that, perhaps even more than *Euryceros*, have beaks in which the upper and lower bill appear unusually decoupled.

In addition to the small size of the clade, a potential shortcoming of these analyses is that the methods assume an underlying Brownian model of evolution, which we have shown is a poor fit for our shape data. Although alternative models have not been developed, one proposed solution to partially account for this is using rescaled phylogenies that better reflect rates of trait evolution in the clade (Zelditch and Goswami 2021; Larouche et al. 2023).

### Rapid morphological evolution, but little evidence for an early burst

The net rate of evolution was 2-4 times higher in the Malagasy than the non-Malagasy vangas for most ecomorphological traits, but the best supported models in our variables rates analyses did not include an early burst coincident with the colonization of Madagascar. Instead, our results support a model of high but uneven rates of recent trait diversification, with many individual taxa diverging dramatically and in different directions. In the skull these shifts are concentrated in the subclade of Malagasy vangas with derived (probing and sallying) foraging behaviors, which also show an overall increase in rates of size evolution. In contrast, foot shape evolutionary rates have generally increased through time across all vangas, with a single dramatic increase in *Hypositta*.

Our dtt results also do not support an early burst pattern. In the classic model of an early burst, we expect disparity to partition between subclades early in the radiation’s history, resulting in a rapid drop in relative subclade disparity. Instead, subclade disparity for skull shape was initially within the null distribution, then increased and stayed well above BM expectations during the diversification of the most morphologically disparate sallying and probing vangas. Foot shape disparity never departed significantly from BM. Taken together, these results show that most of the morphological diversification of the Malagasy vangas occurred after they had already been present in Madagascar for millions of years and coincident with the evolution of novel foraging behaviors.

The delayed increase in bill shape diversification, limited to a clade in which derived foraging behaviors have evolved, is consistent with the model of a subclade-specific key innovation (Etienne and Haegeman 2012; Slater and Pennell 2014). These major categories of foraging behavior do not correspond directly with the diverse specific behaviors employed by vangas with specialized bill and foot shapes (e.g. gleaning from tree trunks vs. leaf litter on the forest floor) but may have functioned as general behavioral innovations which permitted further diversification. A range of extrinsic or intrinsic factors could have played a role in the timing of this shift. Extrinsic factors include shifts in climatic or competitive dynamics with other clades. One possible intrinsic factor is limited genetic diversity due to small founder populations. A hybrid origin is increasingly recognized as common in adaptive radiations (Seehausen 2004; Marques et al. 2019; Martin and Richards 2019; Gillespie et al. 2020; Wogan et al. 2023), and future work should test for the presence of reticulation events, which were recently shown to have contributed to diversification in gemsnakes, another Malagasy radiation of similar age (DeBaun et al. 2023). Some of these factors could also have contributed to the shifts in substrate use associated with pedal morphology.

The idea that founder niche can constrain adaptive radiation, and that certain intrinsic traits may predispose some lineages to diversify, is also widespread in the literature (Jablonski 2008; Flohr et al. 2013). In birds, granivory has been proposed as a trait which facilitates diversification, largely because the two best-studied avian radiations, the Galapagos finches and Hawaiian honeycreepers, both evolved high dietary diversity from a granivorous ancestor (Lovette et al. 2002b; Rundell and Price 2009). Constraint imposed by insectivorous ancestry is therefore another possible explanation for the delay in bill shape diversification in Vangidae. Although they have evolved numerous means of foraging for insects, and the larger Malagasy vangas also prey on small vertebrates, they have not evolved the dietary diversity of the other classic avian radiations. This restriction might be one explanation for the slower initial diversification rate in vangas. A different, though related explanation is that vangas may belong to a clade with lower baseline evolvability. Passerida, the large clade in which the finches and honeycreepers are both nested, shows exceptionally high rates of both morphological evolution and speciation (Lovette et al. 2002b; Felice and Goswami 2018; Oliveros et al. 2019; Vinciguerra and Burns 2021; Imfeld and Barker 2022), indicating a high propensity for diversification. In contrast, the vangas are in Corvides, which show support for an increase in speciation rate at their base (Oliveros et al. 2019) but no corresponding overall increase in rates of skull shape evolution (Felice and Goswami 2018).

An unavoidable limitation, in the case of Vangidae, is our use of extant-only methods to analyze patterns of diversification over time. Extant-only methods often fail to detect the patterns of diversification through time preserved in the fossil record (Mitchell 2015), and this is likely to be a greater concern for older clades which have had more opportunity for extinction to impact present diversity patterns. Unfortunately the fossil record of birds generally is extremely poor due to low preservation potential, and few, if any, fossils exists in Madagascar between the late Cretaceous and late Pleistocene.

### Integrating multiple traits and outliers

The early burst model of adaptive radiation assumes a clade-wide effect in which diversification is distributed uniformly across species. The Malagasy vangas offer a very different model of adaptive radiation, where instead many individual taxa diverge dramatically in one or more aspects of their morphology to generate most of the clade’s diversity. This presents substantial challenges to the quantitative analysis of trait diversity and evolution-existing conceptual frameworks and empirical tests for understanding adaptive radiation are not well-designed for clades in which massive outliers are essential to understanding the clade as a whole. This is not a new challenge. For example, (Slater et al. 2010; Rowsey et al. 2019) both found early bursts in morphological evolution only after removing outliers from their respective clades. Our confidence in the results of several analyses was complicated due to both the small number of taxa in Vangidae and the large impact of outliers. This challenge may be inherent to our statistical approaches, but nonetheless needs to be accounted for in our biological interpretations.

Our analysis of both bill and foot shape demonstrated that the adaptive radiation of vangas has proceeded along multiple trait axes. These traits show a lack of integration and have very different patterns of disparity, indicating that they have made generally independent contributions to vanga diversity. Different species display extreme morphologies in bill or foot shape, facilitating their specialization to an exceptionally wide range of foraging niches, though *Mystacornis* and *Oriolia* do show some specialization in both bill and pedal morphology. As important as bill shape is to adaptive diversification in birds, studying only that trait would have underestimated the ecomorphological diversity present in the Malagasy vangas and simplified our understanding of evolutionary tempo in the clade.

### Conclusions and Future Directions

Vangas offer an example of an adaptive radiation that has achieved exceptional ecomorphological diversity, but not through an early burst of trait evolution. Our study demonstrated the importance of using multiple traits to examine the complexities of morphological diversification, especially in clades that might be responding to various selective pressures. We emphasized the importance of exceptional ecomorphological diversity as the unifying feature of adaptive radiations, and explored the impact of extreme phenotypes in driving diversification patterns. Our conclusions about integration and modularity were highly influenced by inclusion or removal of the extreme taxa, indicating that there are idiosyncratic patterns of trait covariation in anatomically divergent vangas. These results suggest that integration patterns may be generally conserved but can also evolve relatively rapidly in specific traits under selective pressure. The evolution of novel classes of foraging behavior can be considered a ‘key innovation’ in Malagasy vangas, but this alone does not explain the higher phenotypic diversity in this clade as other related factors such as locomotory mode have driven diversification in different directions.

In this study we took a sister clade approach to evaluating the diversification of Malagasy vangas. However a more complete understanding of biodiversity patterns and their causes will require broadening taxonomic scope to comparisons across a wider range of taxa, without losing the insights that emerge from species-level sampling (Losos and Miles 2002; Moen et al. 2021). We highlight the importance of future studies to explore the frequency of outliers, i.e. species exhibiting extremely divergent morphologies, across traits and taxonomic scales, and to investigate whether this is a more general feature of many adaptive radiations, as some other authors have suggested (Slater and Pennell 2014; Martin and Richards 2019; Rowsey et al. 2020). A survey of this phenomenon would help evaluate how extreme phenotype should fit into our conceptual frameworks and be accounted for in analyses of macroevolution.

## Author contributions

A.A. and S.R. designed research; A.A., E.L. S.R. performed research and analyzed data; A.A. and S.R. wrote the paper with contributions from all authors.

## Acknowledgments

We gratefully acknowledge the collections staff for use of specimens sampled for this study, including Ben Marks, John Bates, and Shannon Hackett at the Field Museum and Paul Sweet and Thomas Trombone at the American Museum of Natural History. We thank Kelsi Hurdle and Aki Watanabe (NYIT), David Birlenbach, Gabriele Ilarde, and Francesca Socki (UMN), and April Neander and Zhe-Xi Luo (UChicago) for CT scanning access and assistance, as well as the use of the scanner at Loyola University Chicago. Access to scans on Morphosource provided by Roger Benson, and the Field and Yale Peabody museums with funding from the oVert project. We thank Kieran McNulty, Keith Barker, Shanta Hejmadi, Sharon Jansa, and Tyler Imfeld for their help and feedback on earlier versions of this manuscript, and then entire Reddy-Barker-Jansa-Ford lab group for helpful discussion and support.

---

**Figure S1.**
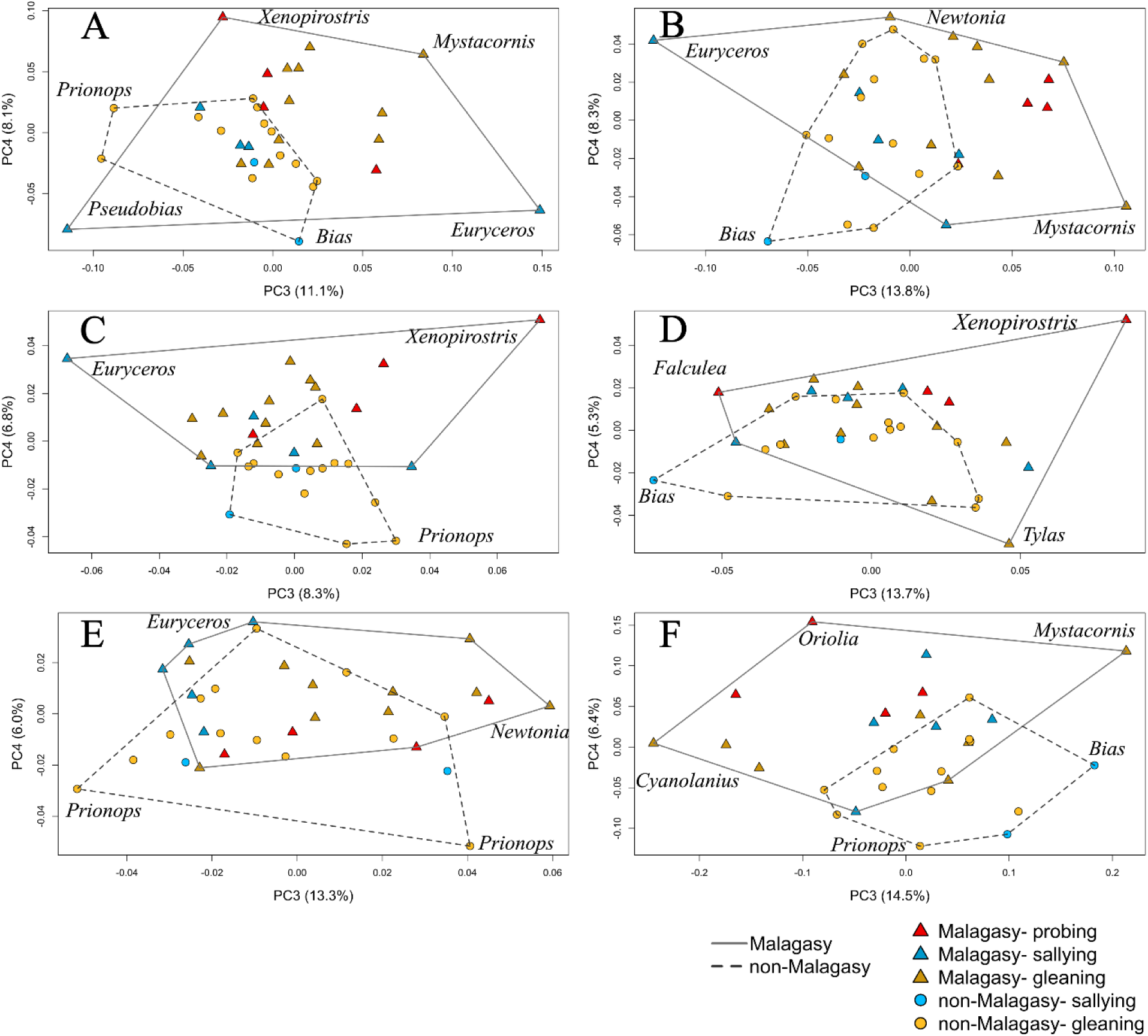
Principal components 3 and 4 of vanga morphological diversity. Top row: A) whole skull and A) whole skull posthoc rotation scores. Middle: C) upper bill and D) lower bill. Bottom: E) cranium and F) feet. Triangles represent Malagasy vangas, while circles represent non-Malagasy vangas. Points are colored by foraging category, with red for probing, blue for sallying, and yellow for gleaning; darker shades are used for the Malagasy vangas. Convex hulls are drawn around each clade, solid (Malagasy) and dashed (non-Malagasy).

**Figure S2.**
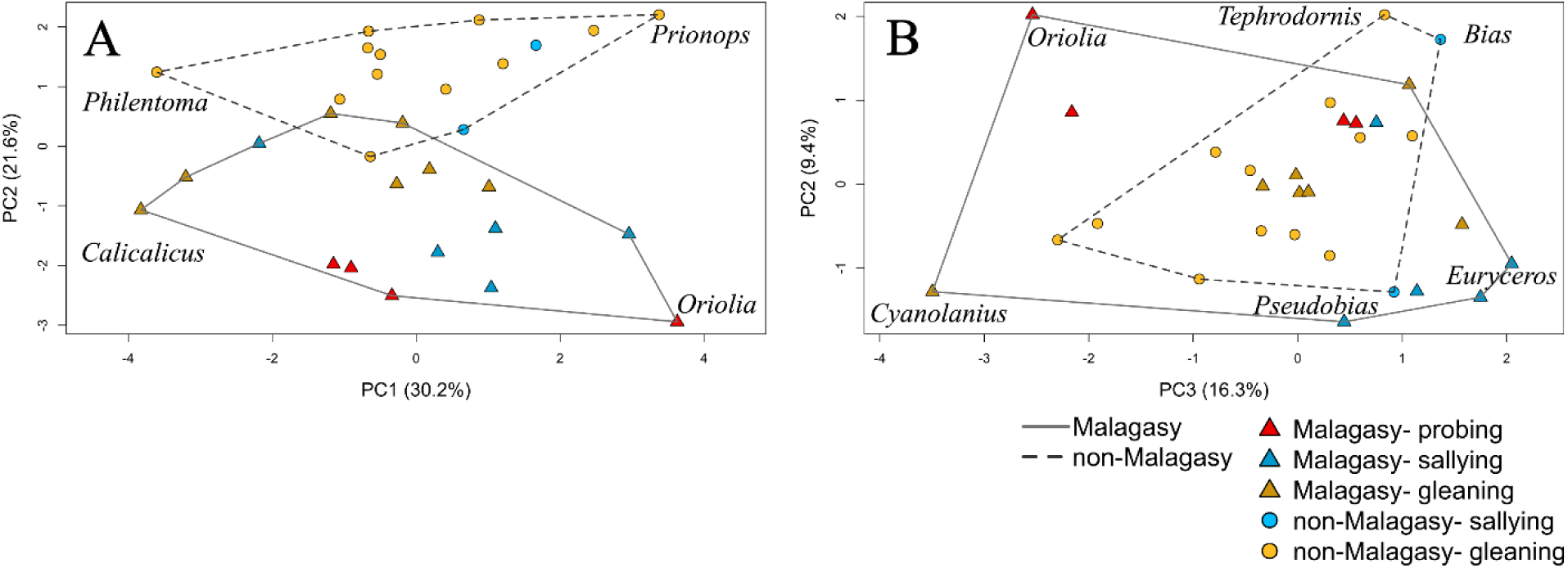
Principal components for the foot dataset with posthoc rotation of outliers. PC scores 1-2 (A) and 3-4 (B). Notably neither *Mystacornis* nor *Hypositta* remain even at the extreme ends of any of these axes, indicating different dominant patterns of variance in the rest of the vangas. Triangles represent Malagasy vangas, while circles represent non-Malagasy vangas. Points are colored by foraging category, with red for probing, blue for sallying, and yellow for gleaning; darker shades are used for the Malagasy vangas. Convex hulls are drawn around each clade, solid (Malagasy) and dashed (non-Malagasy).

**Figure S3.**
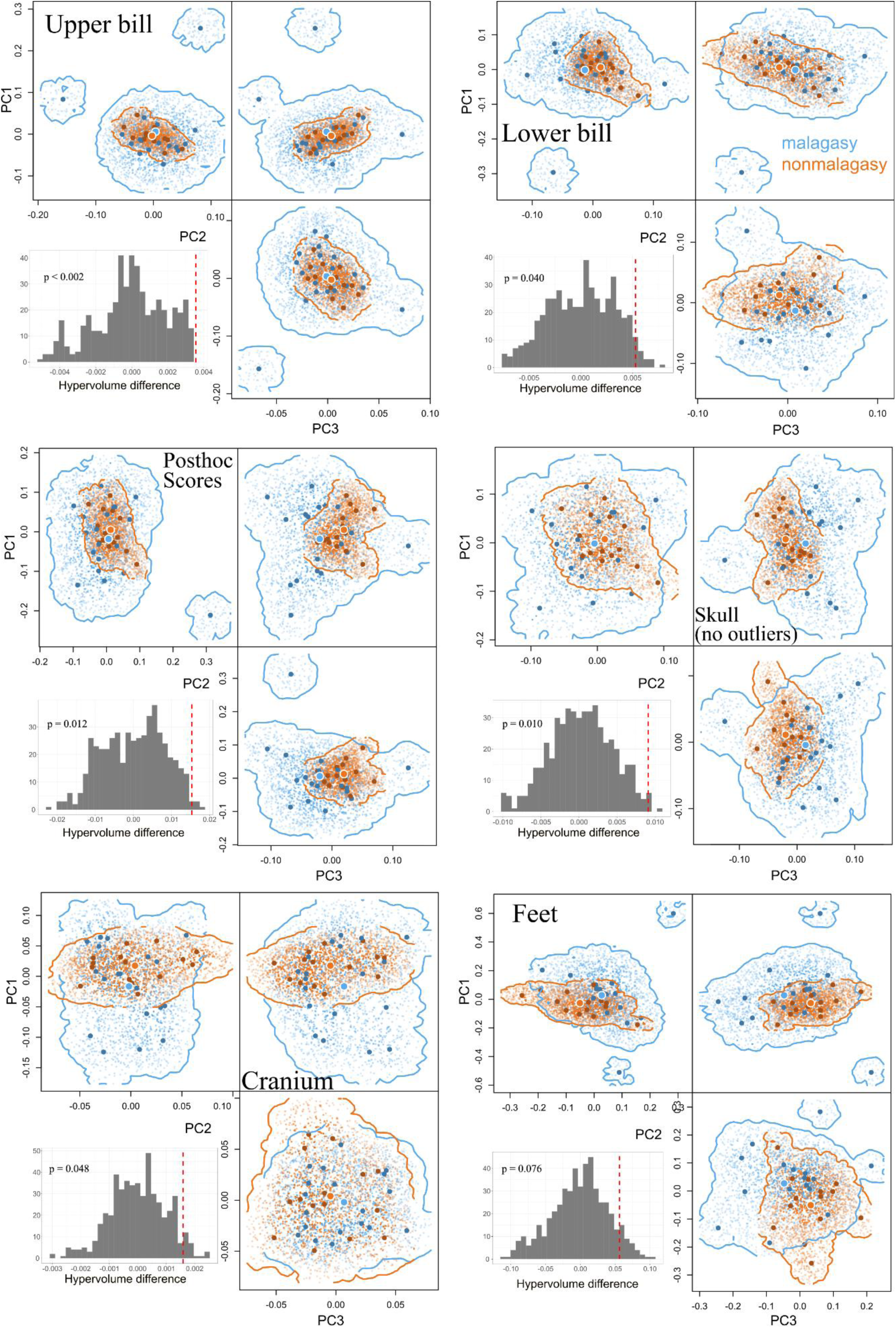
Morphospace hypervolumes by module for Malagasy (blue) and non-Malagasy (orange) vangas. Hypervolumes generated using the first three PC axes for each module. The inset histogram is the null distribution of hypervolume differences from the permutation test, with a dashed line indicating the observed difference in volume. Modules are the upper and lower bill (top row); whole skull (middle row), using posthoc rotation scores (left) or with the two bill shape outliers excluded (right); the cranium (bottom left); and the feet (bottom right). Differences in morphospace occupancy were significantly different for all skull modules, but not for the feet.

**Figure S4.**
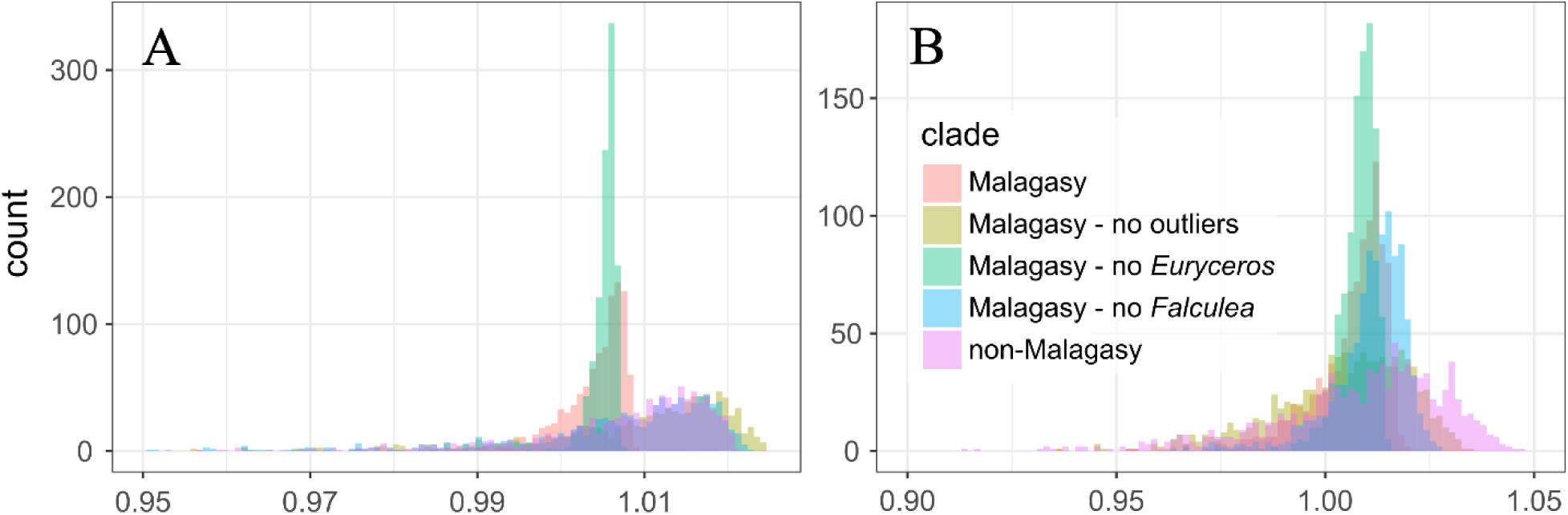
Null distribution of the covariance ratio from the permutation procedure of the modularity test for A) the upper and lower bill and B) the upper bill and skull. The inclusion of anatomical outliers can result in a very different mean and variance for the null distribution (red, green and blue). When outliers are removed, the distribution for the Malagasy and non-Malagasy vangas is similar (yellow and pink). For modularity of the upper and lower bill (A), only *Falculea* seems to contribute to this pattern, while for the upper bill and skull both outliers contribute.

**Figure S5.**
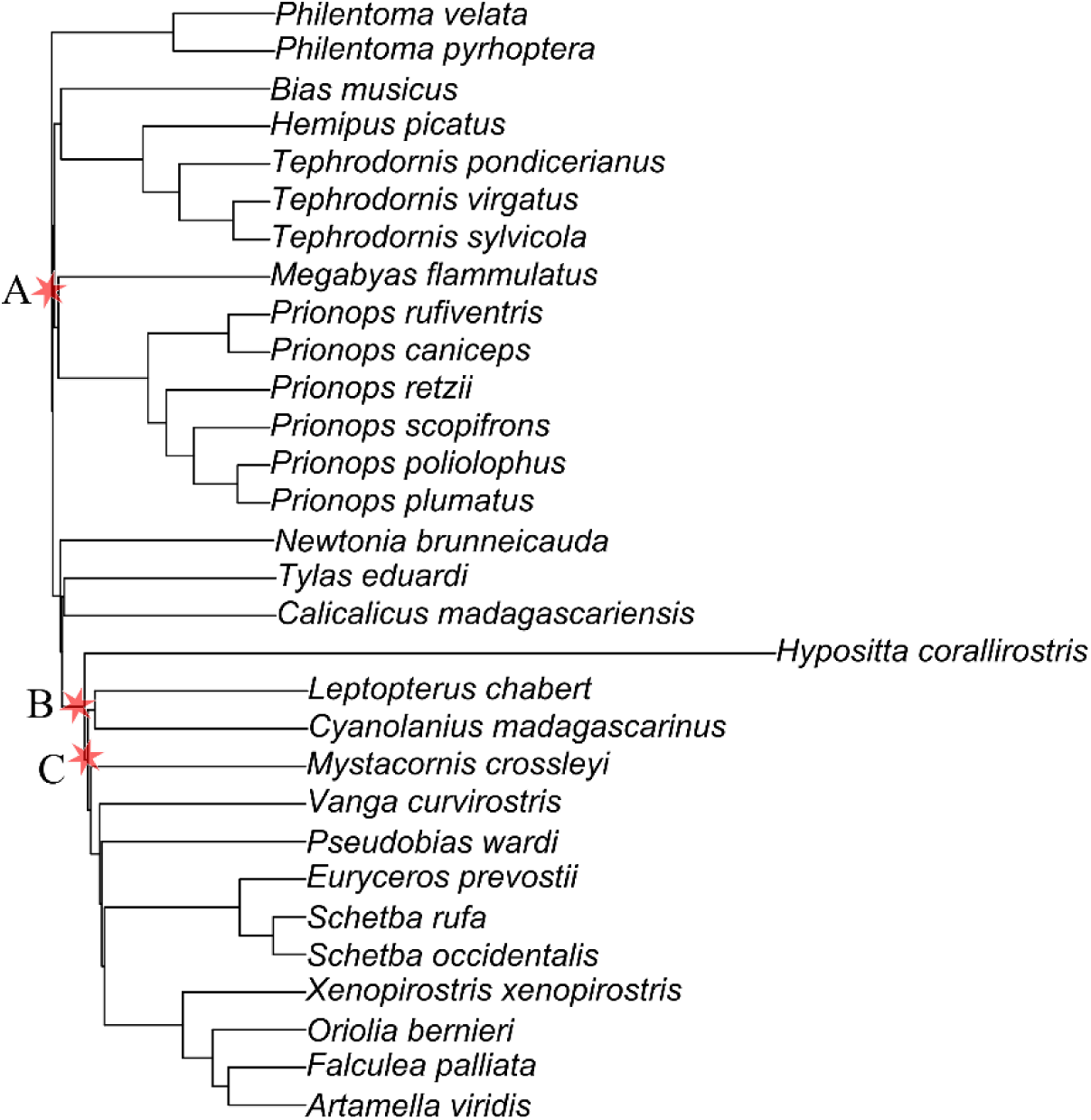
Vanga phylogeny rescaled by mean delta transforms from the best supported model, showing estimated rates of foot shape evolution. Overall, branch lengths are compressed towards the root as rates of evolution increase through time, with a burst of evolution at *Hypositta* (Nuthatch-Vanga). Stars indicate positions of transforms, corresponding to nodes 1 (A), 5 (B) and 6 (C) in Table S10.

**Figure S6:**
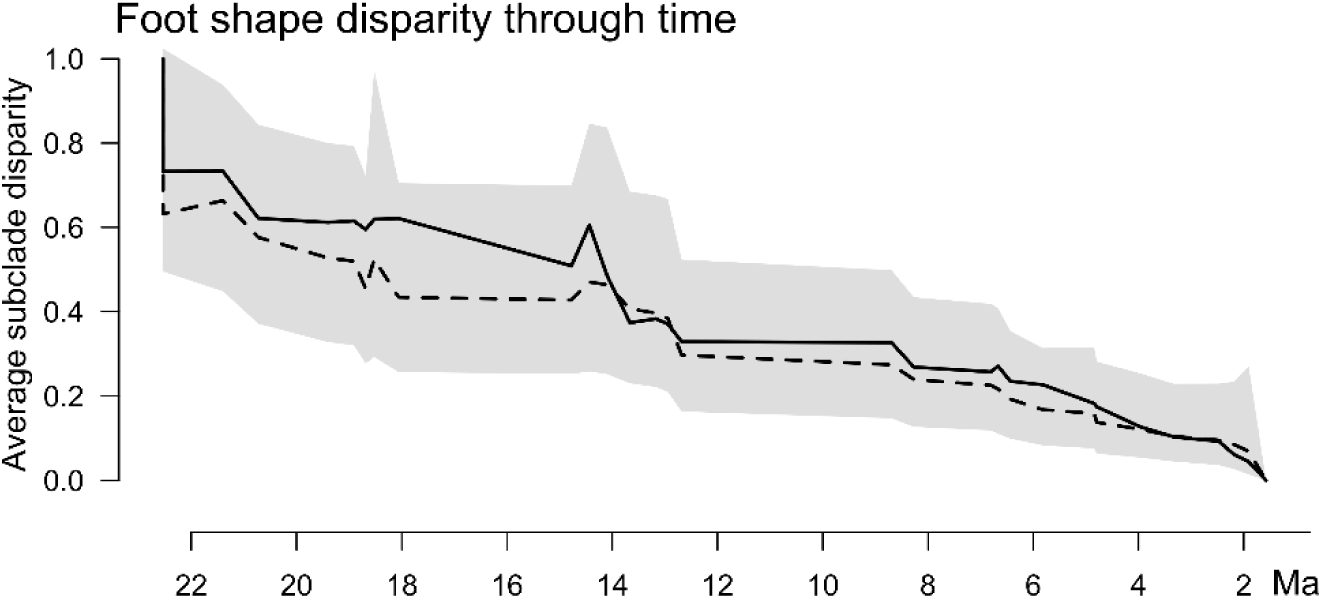
Disparity through time for foot shape, using the PC scores representing 95% of variation in foot shape. Solid line indicates observed disparity, dashed line simulated disparity under constant rates, and the shaded area the 95% confidence interval. Overall subclade disparity in foot shape does not deviate from expectations under Brownian motion.

**Table S1.**
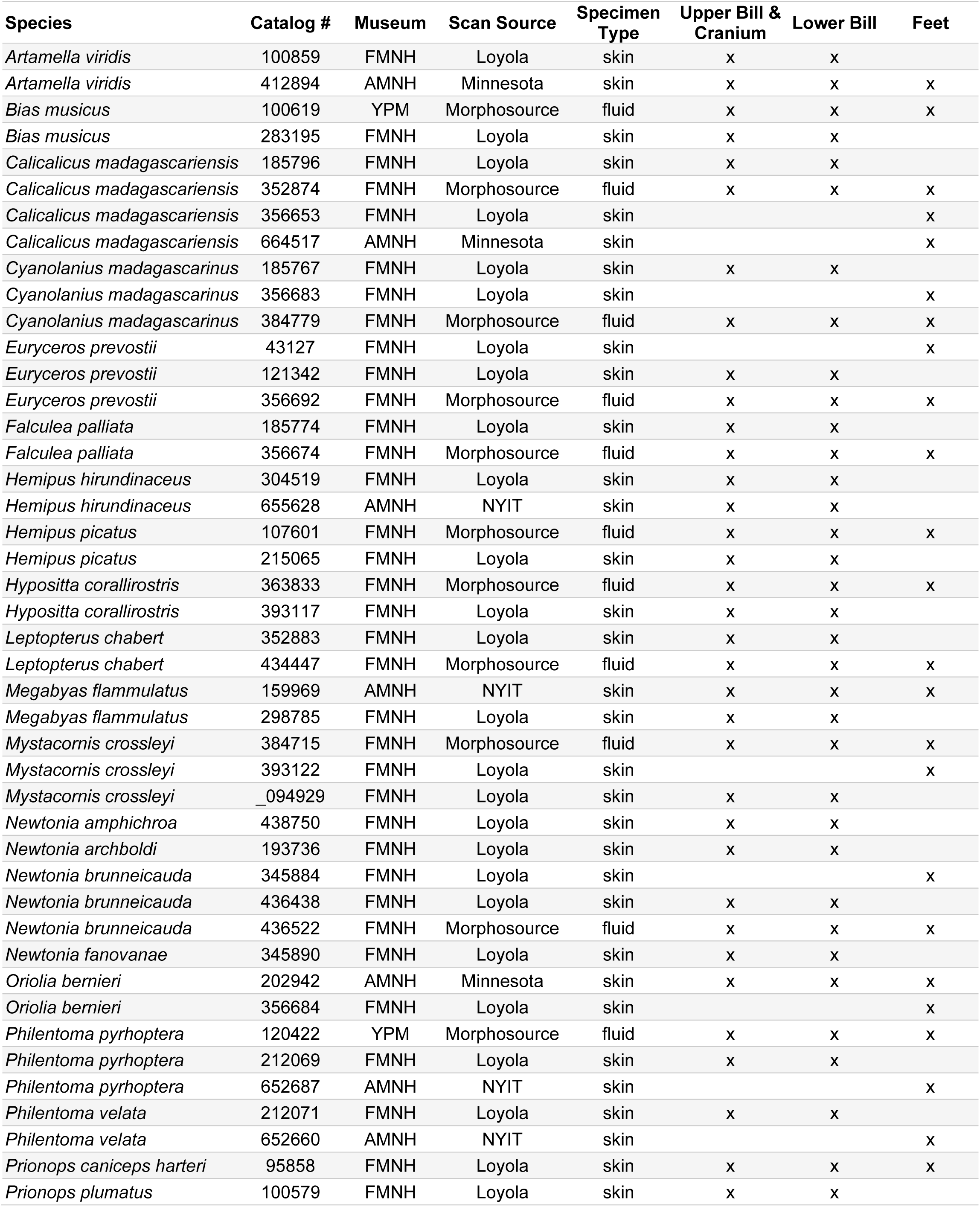
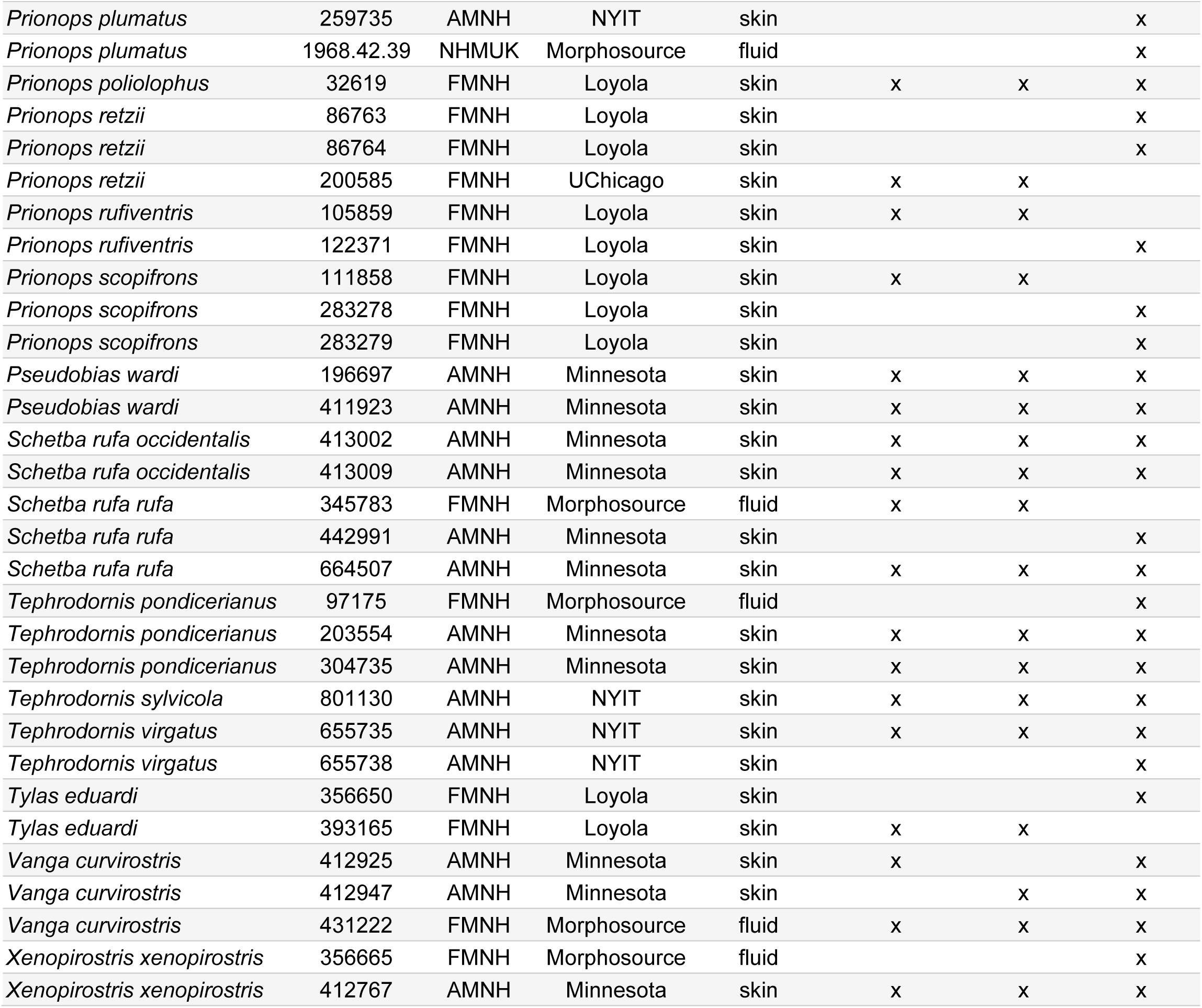
Specimen and scan information for the 75 museum specimens included in this study. Scan source specifies the institution where the specimen was CT scanned or if the scan was downloaded from Morphosource. Specimen type is whether the specimen is a round skin or fluid (alcohol) preserved specimen. We indicate which anatomical regions were measured for each specimen; note that all skull modules were measured on the same specimens except for two specimens of *Vanga curvirostris*, where damage prevented us from doing so. For fluid specimens, missing information typically indicates that parts of the specimen were damaged or missing, but many round skins had only the skull or feet scanned on a particular specimen.

| Species | Catalog # | Museum | Scan Source | Specimen Type | Upper Bill & Cranium | Lower Bill | Feet |
| --- | --- | --- | --- | --- | --- | --- | --- |
| *Artamella viridis* | 100859 | FMNH | Loyola | skin | x | x |  |
| *Artamella viridis* | 412894 | AMNH | Minnesota | skin | x | x | x |
| *Bias musicus* | 100619 | YPM | Morphosource | fluid | x | x | x |
| *Bias musicus* | 283195 | FMNH | Loyola | skin | x | x |  |
| *Calicalicus madagascariensis* | 185796 | FMNH | Loyola | skin | x | x |  |
| *Calicalicus madagascariensis* | 352874 | FMNH | Morphosource | fluid | x | x | x |
| *Calicalicus madagascariensis* | 356653 | FMNH | Loyola | skin |  |  | x |
| *Calicalicus madagascariensis* | 664517 | AMNH | Minnesota | skin |  |  | x |
| *Cyanolanius madagascarinus* | 185767 | FMNH | Loyola | skin | x | x |  |
| *Cyanolanius madagascarinus* | 356683 | FMNH | Loyola | skin |  |  | x |
| *Cyanolanius madagascarinus* | 384779 | FMNH | Morphosource | fluid | x | x | x |
| *Euryceros prevostii* | 43127 | FMNH | Loyola | skin |  |  | x |
| *Euryceros prevostii* | 121342 | FMNH | Loyola | skin | x | x |  |
| *Euryceros prevostii* | 356692 | FMNH | Morphosource | fluid | x | x | x |
| *Falcula palliata* | 185774 | FMNH | Loyola | skin | x | x |  |
| *Falcula palliata* | 356674 | FMNH | Morphosource | fluid | x | x | x |
| *Hemipus hirundinaceus* | 304519 | FMNH | Loyola | skin | x | x |  |
| *Hemipus hirundinaceus* | 655628 | AMNH | NYIT | skin | x | x |  |
| *Hemipus picatus* | 107601 | FMNH | Morphosource | fluid | x | x | x |
| *Hemipus picatus* | 215065 | FMNH | Loyola | skin | x | x |  |
| *Hypositta corallirostris* | 363833 | FMNH | Morphosource | fluid | x | x | x |
| *Hypositta corallirostris* | 393117 | FMNH | Loyola | skin | x | x |  |
| *Leptopterus chabert* | 352883 | FMNH | Loyola | skin | x | x |  |
| *Leptopterus chabert* | 434447 | FMNH | Morphosource | fluid | x | x | x |
| *Megabyas flammulatus* | 159969 | AMNH | NYIT | skin | x | x | x |
| *Megabyas flammulatus* | 298785 | FMNH | Loyola | skin | x | x |  |
| *Mystacornis crossleyi* | 384715 | FMNH | Morphosource | fluid | x | x | x |
| *Mystacornis crossleyi* | 393122 | FMNH | Loyola | skin |  |  | x |
| *Mystacornis crossleyi* | _094929 | FMNH | Loyola | skin | x | x |  |
| *Newtonia amphichroa* | 438750 | FMNH | Loyola | skin | x | x |  |
| *Newtonia archboldi* | 193736 | FMNH | Loyola | skin | x | x |  |
| *Newtonia brunneicauda* | 345884 | FMNH | Loyola | skin |  |  | x |
| *Newtonia brunneicauda* | 436438 | FMNH | Loyola | skin | x | x |  |
| *Newtonia brunneicauda* | 436522 | FMNH | Morphosource | fluid | x | x | x |
| *Newtonia fanovanae* | 345890 | FMNH | Loyola | skin | x | x |  |
| *Oriolia bernieri* | 202942 | AMNH | Minnesota | skin | x | x | x |
| *Oriolia bernieri* | 356684 | FMNH | Loyola | skin |  |  | x |
| *Philentoma pyrhoptera* | 120422 | YPM | Morphosource | fluid | x | x | x |
| *Philentoma pyrhoptera* | 212069 | FMNH | Loyola | skin | x | x |  |
| *Philentoma pyrhoptera* | 652687 | AMNH | NYIT | skin |  |  | x |
| *Philentoma velata* | 212071 | FMNH | Loyola | skin | x | x |  |
| *Philentoma velata* | 652660 | AMNH | NYIT | skin |  |  | x |
| *Prionops caniceps harteri* | 95858 | FMNH | Loyola | skin | x | x | x |
| *Prionops plumatus* | 100579 | FMNH | Loyola | skin | x | x |  |
| *Prionops plumatus* | 259735 | AMNH | NYIT | skin |  |  | x |
| *Prionops plumatus* | 1968.42.39 | NHMHUK | Morphosource | fluid |  |  | x |
| *Prionops poliophus* | 32619 | FMNH | Loyola | skin | x | x | x |
| *Prionops retzii* | 86763 | FMNH | Loyola | skin |  |  | x |
| *Prionops retzii* | 86764 | FMNH | Loyola | skin |  |  | x |
| *Prionops retzii* | 200585 | FMNH | UChicago | skin | x | x |  |
| *Prionops rufiventris* | 105859 | FMNH | Loyola | skin | x | x |  |
| *Prionops rufiventris* | 122371 | FMNH | Loyola | skin |  |  | x |
| *Prionops scopifrons* | 111858 | FMNH | Loyola | skin | x | x |  |
| *Prionops scopifrons* | 283278 | FMNH | Loyola | skin |  |  | x |
| *Prionops scopifrons* | 283279 | FMNH | Loyola | skin |  |  | x |
| *Pseudobias wardi* | 196697 | AMNH | Minnesota | skin | x | x | x |
| *Pseudobias wardi* | 411923 | AMNH | Minnesota | skin | x | x | x |
| *Schetba rufa occidentalis* | 413002 | AMNH | Minnesota | skin | x | x | x |
| *Schetba rufa occidentalis* | 413009 | AMNH | Minnesota | skin | x | x | x |
| *Schetba rufa rufa* | 345783 | FMNH | Morphosource | fluid | x | x |  |
| *Schetba rufa rufa* | 442991 | AMNH | Minnesota | skin |  |  | x |
| *Schetba rufa rufa* | 664507 | AMNH | Minnesota | skin | x | x | x |
| *Tephrodornis pondicerianus* | 97175 | FMNH | Morphosource | fluid |  |  | x |
| *Tephrodornis pondicerianus* | 203554 | AMNH | Minnesota | skin | x | x | x |
| *Tephrodornis pondicerianus* | 304735 | AMNH | Minnesota | skin | x | x | x |
| *Tephrodornis sylvicola* | 801130 | AMNH | NYIT | skin | x | x | x |
| *Tephrodornis virgatus* | 655735 | AMNH | NYIT | skin | x | x | x |
| *Tephrodornis virgatus* | 655738 | AMNH | NYIT | skin |  |  | x |
| *Tylas eduardi* | 356650 | FMNH | Loyola | skin |  |  | x |
| *Tylas eduardi* | 393165 | FMNH | Loyola | skin | x | x |  |
| *Vanga curvirostris* | 412925 | AMNH | Minnesota | skin | x |  | x |
| *Vanga curvirostris* | 412947 | AMNH | Minnesota | skin |  | x | x |
| *Vanga curvirostris* | 431222 | FMNH | Morphosource | fluid | x | x | x |
| *Xenopirostris xenopirostris* | 356665 | FMNH | Morphosource | fluid |  |  | x |
| *Xenopirostris xenopirostris* | 412767 | AMNH | Minnesota | skin | x | x | x |

**Table S2.** Descriptions of all landmarks, semilandmark curves, and linear measurements used for morphometric data in this study. Measurement name refers to the name used in our data files. Anatomical descriptions made with reference to: (Baumel 1993; Felice and Goswami 2018; Navalón et al. 2020; Gómez and Lois-Milevicich 2024).

| Module | Measurement name | Type | Definition |
| --- | --- | --- | --- |
| Cranium | upper_base_med | landmark | midpoint of the craniofacial hinge |
| Cranium | orbit_ant | landmark | lateralmost point of the lacrimal |
| Cranium | orbit_post | landmark | postorbital process, anterior tip |
| Cranium | cranium_med | landmark | posterior end of the frontal depression, medial to the posteriomedial point of the orbit |
| Cranium | orbit | semilandmark curve | 25 landmarks along the edge of the orbit, from orbit_ant to orbit_post |
| Cranium | cranium | semilandmark curve | 10 landmarks along the midline of the cranium from upper_base_med to cranium_med |
| Upper bill | upper_tip | landmark | tip of maxilla, midline, such that the landmark points directly downwards |
| Upper bill | upper_curve | landmark | dorsalmost inflection point/maximum curvature of the tomial edge |
| Upper bill | upper_nasal_base | landmark | proximal edge of the ventral end of the nasal bar projected down to the tomium |
| Upper bill | upper_nares_prox | landmark | posteriormost point of the nasal opening |
| Upper bill | upper_nares_dist | landmark | anteriormost point of the nasal opening |
| Upper bill | max_underside_base | landmark | posteriormost point of the maxilla, medial to join with maxillary processes of palatine |
| Upper bill | maxilla_midline | semilandmark curve | 22 landmarks along the midline of the dorsal surface of the maxilla |
| Upper bill | nares | semilandmark curve | 22 landmarks circling the nares |
| Upper bill | maxmid_under | semilandmark curve | 22 landmarks along the midline of the ventral (inner) surface of the maxilla |
| Upper bill | tomium | semilandmark curve | 25 landmarks along the tomium (cutting edge) of the maxilla |
| Lower bill | lower_tip | landmark | tip of the mandible, midline |
| Lower bill | lowertom_end | landmark | angle of the mandible |
| Lower bill | lower_base | landmark | mandibular symphysis, posterior midline |
| Lower bill | lower_tomium | semilandmark curve | 25 landmarks along the tomium (cutting edge) of the mandible |
| Lower bill | lower_inner | semilandmark curve | 22 landmarks along the midline of the dorsal (inner) surface of the mandible |
| Lower bill | lower_outside | semilandmark curve | 28 landmarks along midline of the ventral (outer) surface of mandible |
| Foot | tarsometatarsus (tmt) | linear measurement | center of the ridge just below the eminentia intercondylaris to distalmost point on the trochlea |
| Foot | metatarsus (m1) | linear measurement | proximal fusion point to tmt to center of the area intercondylaris |
| Foot | hallux (d1p1) | linear measurement | between highest points of the ridges below and above the proximal and distal eminentia intercondylaris |
| Foot | digit 2 phalange 1 (d2p1) | linear measurement | apex of the proximal crest to the center of the divot at the distal end of the bone |
| Foot | d2p2 | linear measurement | proximal to distal end along the midline |
| Foot | d3p1 | linear measurement | proximal to distal end along the midline |
| Foot | d3p2 | linear measurement | proximal to distal end along the midline |
| Foot | d3p3 | linear measurement | proximal to distal end along the midline |
| Foot | d4p1 | linear measurement | proximal to distal end along the midline |
| Foot | d4p2 | linear measurement | proximal to distal end along the midline |
| Foot | d4p3 | linear measurement | proximal to distal end along the midline |
| Foot | d4p4 | linear measurement | proximal to distal end along the midline |

**Table S3.** Shape differences (ANOVA) by pedal bone between clades and foraging groups. Overall differences in foot proportions between Malagasy and non-Malagasy vangas are spread across the first three digits (d1-d3), while differences in shape between foraging groups are in the metacarpal (m1) and digit 3 (the central forward-facing toe). Numbers are p values from each ANOVA.

| Bone | Clades (Malagasy vs non-Mal) |  | Foraging categories |  |
| --- | --- | --- | --- | --- |
|  | phylogenetic | non-Phylo | phylogenetic | non-Phylo |
| tmt | 0.347 | 0.441 | 0.351 | 0.202 |
| m1 | 0.945 | 0.889 | 0.073 | **0.001** |
| d1p1 | 0.561 | **0.026** | 0.943 | 0.444 |
| d2p1 | 0.433 | **0.044** | 0.826 | 0.505 |
| d2p2 | 0.161 | **0.011** | 0.542 | 0.221 |
| d3p1 | 0.199 | **0.002** | 0.113 | **0.001** |
| d3p2 | 0.525 | **0.029** | 0.846 | 0.249 |
| d3p3 | 0.487 | 0.636 | 0.737 | 0.503 |
| d4p1 | 0.516 | 0.295 | 0.823 | 0.467 |
| d4p2 | 0.825 | 0.677 | 0.878 | 0.646 |
| d4p3 | 0.886 | 0.986 | 0.430 | 0.136 |
| d4p4 | 0.704 | 0.808 | 0.300 | 0.098 |

**Table S4.** Differences in morphological diversity between foraging groups, quantified using disparity and MANOVAs from the full landmark or linear measurement dataset. Numbers are significance of difference (p values) between foraging groups (probing, sallying, or gleaning).

| Module | MANOVAs |  | Disparity |  |  |
| --- | --- | --- | --- | --- | --- |
|  | Phylogenetic | non-Phylo | Probe vs. Glean | Probe vs. Sally | Glean vs. Sally |
| Upper bill | 0.319 | **0.011** | **0.015** | 0.106 | 0.609 |
| Lower bill | 0.265 | **0.018** | **0.036** | 0.078 | 0.769 |
| Cranium | 0.221 | **0.001** | **0.001** | **0.003** | 0.721 |
| Whole skull | 0.272 | **0.001** | **0.012** | **0.042** | 0.645 |
| Skull (no outliers) | 0.953 | **0.001** | **0.013** | **0.028** | 0.930 |
| Feet | 0.705 | 0.069 | 0.646 | 0.753 | 0.342 |
| Feet (no outliers) | 0.372 | **0.001** | 0.520 | 0.501 | 0.870 |

**Table S5.** Comparisons of the strength of phylogenetic signal between clades. Nearly all modules showed significant phylogenetic signal in all clades, and differences in the degree of phylogenetic signal were nearly all statistically significant. Z scores indicate the strength of phylogenetic signal for that clade and anatomical region, while the remaining numbers are p values indicating whether differences were significant.

| **Upper Bill** | **Malagasy** | **non-Malagasy** | **Malagasy (no outliers)** | **Malagasy (no *Euryceros*)** | **Malagasy (no *Falcullea*)** |
| --- | --- | --- | --- | --- | --- |
| Z<sub>physignal</sub> | 0.044 | **4.000** | **3.491** | NaN | NaN |
| Malagasy | - |  |  |  |  |
| non-Malagasy | **<0.001** | - |  |  |  |
| Malagasy (no outliers) | **<0.001** | **<0.001** | - |  |  |
| Malagasy (no *Euryceros*) | 1.000 | **<0.001** | **<0.001** | - |  |
| Malagasy (no *Falcullea*) | 1.000 | **<0.001** | **<0.001** | 1.000 | - |
| **Lower Bill** |  |  |  |  |  |
| Z<sub>physignal</sub> | **2.667** | **2.785** | **2.476** | **2.754** | **3.062** |
| Malagasy | - |  |  |  |  |
| non-Malagasy | **0.008** | - |  |  |  |
| Malagasy (no outliers) | **0.008** | **0.005** | - |  |  |
| Malagasy (no *Euryceros*) | **0.006** | **0.006** | **0.006** | - |  |
| Malagasy (no *Falcullea*) | **0.008** | **0.002** | **0.002** | **0.006** | - |
| **Cranium** |  |  |  |  |  |
| Z<sub>physignal</sub> | **4.315** | **2.997** | **4.104** | **4.679** | **4.180** |
| Malagasy | - |  |  |  |  |
| non-Malagasy | **<0.001** | - |  |  |  |
| Malagasy (no outliers) | **<0.001** | **<0.001** | - |  |  |
| Malagasy (no *Euryceros*) | **<0.001** | **<0.001** | **<0.001** | - |  |
| Malagasy (no *Falcullea*) | **<0.001** | **<0.001** | **<0.001** | **0.010** | - |
| **Whole Skull** |  |  |  |  |  |
| Z<sub>physignal</sub> | **3.859** | **3.972** | **5.065** | **4.296** | **3.950** |
| Malagasy | - |  |  |  |  |
| non-Malagasy | **<0.001** | - |  |  |  |
| Malagasy (no outliers) | **<0.001** | **<0.001** | - |  |  |
| Malagasy (no *Euryceros*) | **<0.001** | **<0.001** | **<0.001** | - |  |
| Malagasy (no *Falcullea*) | **<0.001** | **<0.001** | **<0.001** | 0.909 | - |
| **Feet** |  |  |  |  |  |
| Z<sub>physignal</sub> | **3.772** | **4.180** | **3.596** |  |  |
| Malagasy | - |  |  |  |  |
| non-Malagasy | 0.398 | - |  |  |  |
| Malagasy (no outliers) | **<0.001** | **<0.001** | - |  |  |

**Table S6.** Comparisons of within-module integration (eigenvalue dispersion) between clades. The degree to which each module is integrated with itself is quantified using eigenvalue dispersion. Pairwise comparisons of integration found few significant differences between clades. Z score for each pairwise comparison is reported above and p value below the diagonal, with significant differences bolded.

| Upper Bill | Malagasy | non-Malagasy | Malagasy (no outliers) | Malagasy (no Euryceros) | Malagasy (no Falculea) |
| --- | --- | --- | --- | --- | --- |
| Z<sub>Vrel</sub> | -0.230 | -0.672 | -0.870 | 0.137 | -0.219 |
| Malagasy | - | 1.156 | 1.749 | 1.023 | 0.033 |
| non-Malagasy | 0.248 | - | 0.505 | **2.088** | 1.170 |
| Malagasy (no outliers) | 0.080 | 0.614 | - | **2.711** | 1.754 |
| Malagasy (no *Euryceros*) | 0.306 | **0.037** | **0.007** | - | 0.974 |
| Malagasy (no *Falculea*) | 0.974 | 0.242 | 0.079 | 0.330 | - |
| **Lower Bill** |  |  |  |  |  |
| Z<sub>Vrel</sub> | -0.050 | -0.262 | -0.629 | 0.081 | -0.552 |
| Malagasy | - | 0.557 | 1.584 | 0.363 | 1.396 |
| non-Malagasy | 0.578 | - | 0.933 | 0.886 | 0.747 |
| Malagasy (no outliers) | 0.113 | 0.351 | - | 1.911 | 0.209 |
| Malagasy (no *Euryceros*) | 0.716 | 0.376 | 0.056 | - | 1.732 |
| Malagasy (no *Falculea*) | 0.163 | 0.455 | 0.834 | 0.083 | - |
| **Cranium** |  |  |  |  |  |
| Z<sub>Vrel</sub> | -0.494 | -0.860 | -0.711 | -0.676 | -0.493 |
| Malagasy | - | 0.958 | 0.592 | 0.504 | 0.005 |
| non-Malagasy | 0.338 | - | 0.380 | 0.477 | 0.949 |
| Malagasy (no outliers) | 0.554 | 0.704 | - | 0.095 | 0.588 |
| Malagasy (no *Euryceros*) | 0.614 | 0.633 | 0.924 | - | 0.501 |
| Malagasy (no *Falculea*) | 0.996 | 0.342 | 0.557 | 0.616 | - |
| **Whole Skull** |  |  |  |  |  |
| Z<sub>Vrel</sub> | -0.421 | -0.990 | -1.100 | -0.163 | -0.775 |
| Malagasy | - | 1.491 | 1.857 | 0.717 | 0.986 |
| non-Malagasy | 0.136 | - | 0.280 | **2.135** | 0.555 |
| Malagasy (no outliers) | 0.063 | 0.780 | - | **2.521** | 0.874 |
| Malagasy (no *Euryceros*) | 0.474 | **0.033** | **0.012** | - | 1.676 |
| Malagasy (no *Falculea*) | 0.324 | 0.579 | 0.382 | 0.094 | - |
| **Feet** |  |  |  |  |  |
| Z<sub>Vrel</sub> | -0.476 | -0.899 | -0.515 |  |  |
| Malagasy | - | 1.032 | 0.097 |  |  |
| non-Malagasy | 0.302 | - | 0.899 |  |  |
| Malagasy (no outliers) | 0.923 | 0.369 | - |  |  |

**Table S7.** Comparisons of between-module integration between clades. Between-module integration quantified using partial least squares analysis, measuring the strength of covariation between pairs of modules. Pairs of skull modules were sometimes but not always significantly integrated with one another, with the inclusion of outliers having a large impact on these results; significant Z-scores are bolded. Pairwise comparisons of integration found few significant differences between clades. Z scores for each pairwise comparison are reported above and p values below the diagonal, with significant differences bolded.

| **Upper Bill vs Cranium** | **Malagasy** | **non-Malagasy** | **Malagasy (no outliers)** | **Malagasy (no Falculea)** | **Malagasy (no Euryceros)** |
| --- | --- | --- | --- | --- | --- |
| Z<sub>Integration</sub> | **2.417** | 1.737 | 1.276 | **2.562** | 1.196 |
| Malagasy | - | 1.352 | 1.616 | 0.652 | 0.898 |
| non-Malagasy | 0.176 | - | 0.388 | 1.790 | 0.244 |
| Malagasy (no outliers) | 0.106 | 0.698 | - | **1.991** | 0.500 |
| Malagasy (no Falculea) | 0.514 | 0.073 | **0.047** | - | 1.401 |
| Malagasy (no Euryceros) | 0.369 | 0.807 | 0.617 | 0.161 | - |
| **Upper Bill vs Lower Bill** |  |  |  |  |  |
| Z<sub>Integration</sub> | **2.108** | **2.513** | **3.623** | 1.056 | **2.202** |
| Malagasy | - | 1.518 | 1.593 | 1.489 | 0.497 |
| non-Malagasy | 0.129 | - | 0.206 | 0.156 | 1.763 |
| Malagasy (no outliers) | 0.111 | 0.837 | - | 0.053 | 1.817 |
| Malagasy (no Falculea) | 0.137 | 0.876 | 0.958 | - | 1.749 |
| Malagasy (no Euryceros) | 0.619 | 0.078 | 0.069 | 0.080 | - |

**Table S8.** Tests of integration (partial least squares) between feet and skull modules. We found no instances of significant integration between these disparate anatomical regions, although integration of the upper bill with the feet approached significance (p = 0.063) in the non-Malagasy vangas.

| Skull module | *p* value | Z score |
| --- | --- | --- |
| Whole skull | 0.626 | -0.383 |
| Whole skull - Malagasy | 0.539 | -0.189 |
| Whole skull – Malagasy no outliers | 0.599 | -0.238 |
| Whole skull – Malagasy no Euryceros | 0.430 | 0.099 |
| Whole skull – Malagasy no Falculea | 0.740 | -0.704 |
| Whole skull – non- Malagasy | 0.246 | 0.719 |
| Lower bill | 0.425 | 0.162 |
| Lower bill - Malagasy | 0.422 | 0.173 |
| Lower bill – Malagasy no outliers | 0.358 | 0.368 |
| Lower bill – Malagasy no Euryceros | 0.398 | 0.214 |
| Lower bill – Malagasy no Falculea | 0.147 | 1.084 |
| Lower bill – non-Malagasy | 0.637 | -0.368 |
| Cranium | 0.766 | -0.745 |
| Cranium - Malagasy | 0.266 | 0.660 |
| Cranium – Malagasy no outliers | 0.233 | 0.738 |
| Cranium – Malagasy no Euryceros | 0.492 | 0.018 |
| Cranium – Malagasy no Falculea | 0.144 | 1.085 |
| Cranium – non-Malagasy | 0.227 | 0.790 |
| Upper bill | 0.460 | 0.047 |
| Upper bill - Malagasy | 0.472 | 0.052 |
| Upper bill – Malagasy no outliers | 0.338 | 0.487 |
| Upper bill – Malagasy no Euryceros | 0.371 | 0.284 |
| Upper bill – Malagasy no Falculea | 0.467 | 0.035 |
| Upper bill – non-Malagasy | 0.063 | 1.536 |

**Table S9.**
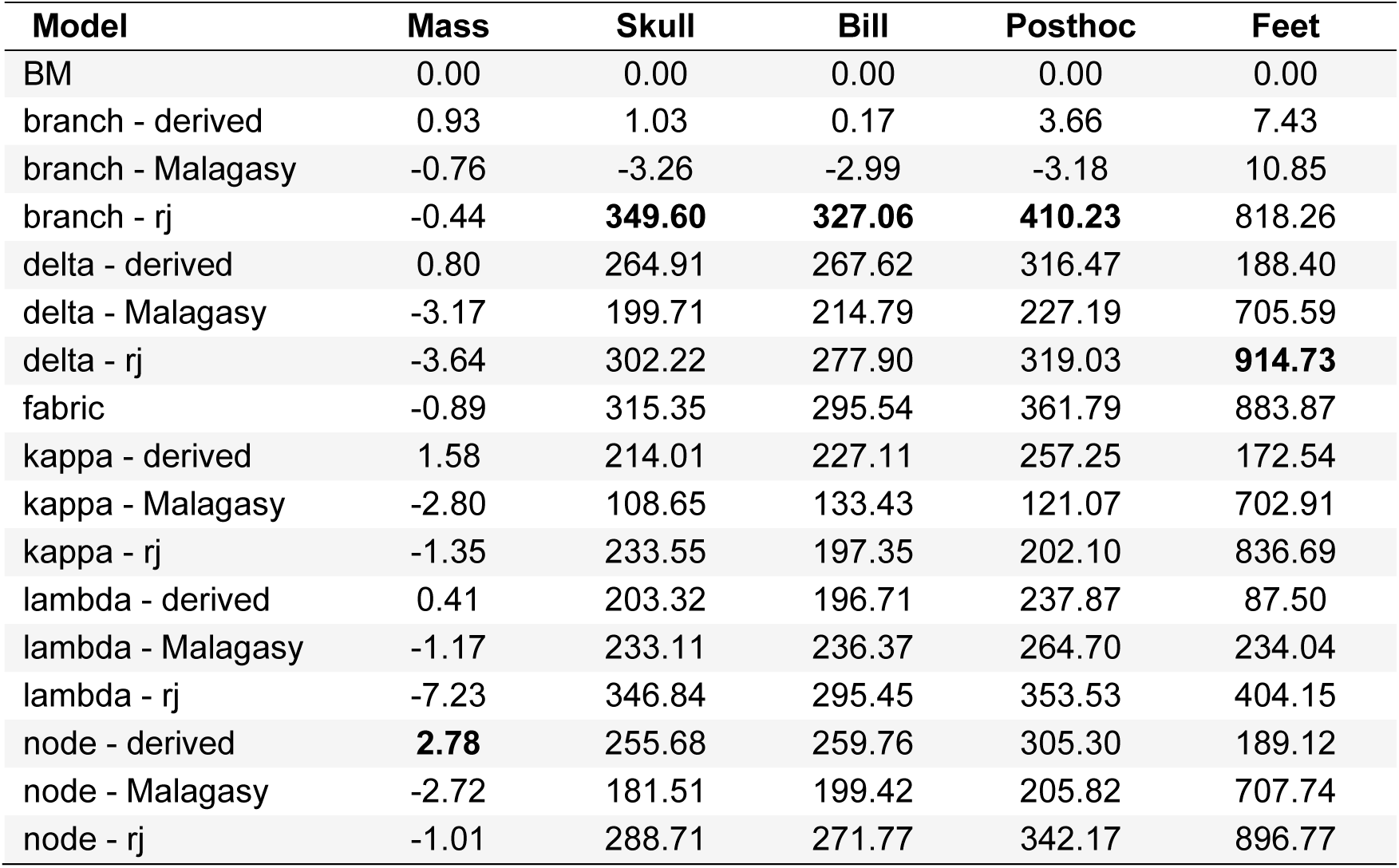
Model support (log Bayes factors) by anatomical region from evolutionary rate (Bayestraits) analyses. Model support for each model was calculated relative to Brownian motion (BM). The best-supported model for each dataset is bolded. Support is averaged across 5 independent runs for each model. Note for body mass only one model received moderate support over BM, whereas for all multivariate shape datasets most models were favored over BM.

| Model | Mass | Skull | Bill | Posthoc | Feet |
| --- | --- | --- | --- | --- | --- |
| BM | 0.00 | 0.00 | 0.00 | 0.00 | 0.00 |
| branch - derived | 0.93 | 1.03 | 0.17 | 3.66 | 7.43 |
| branch - Malagasy | -0.76 | -3.26 | -2.99 | -3.18 | 10.85 |
| branch - rj | -0.44 | **349.60** | **327.06** | **410.23** | 818.26 |
| delta - derived | 0.80 | 264.91 | 267.62 | 316.47 | 188.40 |
| delta - Malagasy | -3.17 | 199.71 | 214.79 | 227.19 | 705.59 |
| delta - rj | -3.64 | 302.22 | 277.90 | 319.03 | **914.73** |
| fabric | -0.89 | 315.35 | 295.54 | 361.79 | 883.87 |
| kappa - derived | 1.58 | 214.01 | 227.11 | 257.25 | 172.54 |
| kappa - Malagasy | -2.80 | 108.65 | 133.43 | 121.07 | 702.91 |
| kappa - rj | -1.35 | 233.55 | 197.35 | 202.10 | 836.69 |
| lambda - derived | 0.41 | 203.32 | 196.71 | 237.87 | 87.50 |
| lambda - Malagasy | -1.17 | 233.11 | 236.37 | 264.70 | 234.04 |
| lambda - rj | -7.23 | 346.84 | 295.45 | 353.53 | 404.15 |
| node - derived | **2.78** | 255.68 | 259.76 | 305.30 | 189.12 |
| node - Malagasy | -2.72 | 181.51 | 199.42 | 205.82 | 707.74 |
| node - rj | -1.01 | 288.71 | 271.77 | 342.17 | 896.77 |

**Table S10.** Frequency and magnitude of transforms in the best supported Bayestraits models. Node number is the numbering of nodes in the phylogeny; the taxon identity of these nodes is indicated for the best-supported shifts. Node numbers are different for the foot dataset because there are fewer taxa represented. Number of scalars indicates the frequency of that scalar (branch or delta transform) in the posterior distribution of 10,000 trees, with 10,000 meaning it was found 100% of the time; scalars detected >75% of the time are bolded. Mean rate is the magnitude of the directional shift for the branch transform, so large values indicate a larger shift in the direction of trait evolution. All highly supported branch scalars were on terminal branches. For the feet, transform indicates the magnitude of the delta transform applied to that node in the best-supported model. Node 5 (*Hypositta*) refers to the node that is the most recent common ancestor of *Hypositta* and all other Malagasy vangas, while Node 6 refers to its immediate descendant, the common ancestor of all Malagasy vangas excluding *Hypositta*.

| Node # (skull) | Skull - branch |  | Posthoc - branch |  | Bill - branch |  | Feet - delta |  |  |
| --- | --- | --- | --- | --- | --- | --- | --- | --- | --- |
|  | # scalars | mean rate | # scalars | mean rate | # scalars | mean rate | Node # (feet) | # scalars | transform |
| 15 (*Falcullea*) | **10000** | **30.306** | **10000** | **40.225** | **10000** | **34.872** | 1 (root) | **10000** | **1.762369** |
| 22 (*Euryceros*) | **10000** | **19.628** | **10000** | **22.717** | **10000** | **22.086** | 5 (*Hypositta*) | 4830 | 1.175442 |
| 52 (*P. rufiventris*) | **9541** | **7.373** | **8836** | **5.737** | 1145 | 1.452 | 6 (w/out *Hypositta*) | 4211 | 0.901766 |
| 46 (*P. plumatus*) | **9332** | **11.169** | **9793** | **12.357** | **9374** | **11.327** | 3 | 450 | 1.0012 |
| 17 (*Xenopirostris*) | **8641** | **4.961** | **9781** | **6.252** | **9717** | **7.638** | 4 | 403 | 1.001883 |
| 37 | 6966 | 4.207 | 1455 | 1.389 | 869 | 1.284 | 34 | 354 | 0.998404 |
| 47 | 6815 | 6.956 | 5740 | 5.109 | 972 | 1.692 | 2 | 327 | 0.998682 |
| 25 | 6828 | 3.251 | 3869 | 1.995 | 3365 | 2.209 | 7 | 276 | 0.999733 |
| 62 | 6633 | 5.487 | 4358 | 3.414 | 2161 | 2.293 | 8 | 0 | 1 |
| 58 | 6569 | 4.844 | 6720 | 5.071 | 4307 | 3.371 | 9 | 0 | 1 |
| 49 (*P. retzii*) | 5308 | 2.990 | 2591 | 1.722 | **7860** | **5.273** | 10 | 0 | 1 |
| 63 (*H. picatus*) | 4334 | 3.455 | 6355 | 4.542 | **7665** | **6.417** | 11 | 0 | 1 |
| 59 | 2776 | 2.423 | 3357 | 2.857 | 903 | 1.358 | 12 | 0 | 1 |
| 16 | 2921 | 2.056 | 1924 | 1.597 | 113 | 1.010 | 13 | 0 | 1 |
| 23 | 936 | 1.204 | 485 | 1.084 | 2029 | 1.679 | 14 | 0 | 1 |
| 20 | 968 | 1.311 | 1968 | 1.730 | 146 | 1.021 | 15 | 0 | 1 |
| 21 | 884 | 1.263 | 2916 | 2.146 | 209 | 1.041 | 16 | 0 | 1 |
| 45 | 791 | 1.342 | 711 | 1.275 | 135 | 1.013 | 17 | 0 | 1 |
| 2 | 552 | 1.356 | 535 | 1.337 | 282 | 1.121 | 18 | 0 | 1 |
| 8 | 563 | 1.300 | 679 | 1.381 | 444 | 1.244 | 19 | 0 | 1 |
| 6 | 478 | 1.348 | 449 | 1.300 | 445 | 1.313 | 20 | 0 | 1 |
| 7 | 474 | 1.360 | 434 | 1.294 | 544 | 1.447 | 21 | 0 | 1 |
| 43 | 431 | 1.189 | 413 | 1.166 | 310 | 1.127 | 22 | 0 | 1 |
| 30 | 401 | 1.235 | 331 | 1.163 | 488 | 1.377 | 23 | 0 | 1 |
| 9 | 361 | 1.207 | 383 | 1.220 | 361 | 1.220 | 24 | 0 | 1 |
| 13 | 374 | 1.175 | 503 | 1.300 | 204 | 1.051 | 25 | 0 | 1 |
| 10 | 359 | 1.187 | 347 | 1.169 | 320 | 1.153 | 26 | 0 | 1 |
| 55 | 385 | 0.974 | 474 | 0.967 | 257 | 0.986 | 27 | 0 | 1 |
| 4 | 344 | 1.170 | 276 | 1.098 | 339 | 1.180 | 28 | 0 | 1 |
| 12 | 271 | 1.071 | 582 | 1.283 | 226 | 1.044 | 29 | 0 | 1 |
| 51 | 258 | 1.049 | 238 | 1.041 | 753 | 1.266 | 30 | 0 | 1 |
| 40 | 251 | 1.075 | 237 | 1.063 | 288 | 1.110 | 31 | 0 | 1 |
| 56 | 256 | 1.056 | 229 | 1.048 | 207 | 1.039 | 32 | 0 | 1 |
| 11 | 220 | 1.040 | 198 | 1.034 | 115 | 1.008 | 33 | 0 | 1 |
| 26 | 241 | 1.067 | 316 | 1.123 | 360 | 1.187 | 35 | 0 | 1 |
| 61 | 266 | 1.059 | 223 | 1.042 | 179 | 1.036 | 36 | 0 | 1 |
| 14 | 221 | 1.039 | 447 | 1.111 | 120 | 1.006 | 37 | 0 | 1 |
| 54 | 195 | 1.031 | 167 | 1.018 | 224 | 1.047 | 38 | 0 | 1 |
| 44 | 219 | 1.043 | 204 | 1.028 | 265 | 1.066 | 39 | 0 | 1 |
| 65 | 203 | 0.991 | 280 | 0.985 | 236 | 0.989 | 40 | 0 | 1 |
| 19 | 192 | 1.023 | 167 | 1.018 | 206 | 1.037 | 41 | 0 | 1 |
| 41 | 187 | 1.031 | 197 | 1.029 | 226 | 1.053 | 42 | 0 | 1 |
| 35 | 179 | 1.029 | 159 | 1.018 | 455 | 1.200 | 43 | 0 | 1 |
| 18 | 181 | 0.991 | 140 | 0.994 | 158 | 0.995 | 44 | 0 | 1 |
| 50 | 196 | 1.032 | 162 | 1.021 | 143 | 1.017 | 45 | 0 | 1 |
| 67 | 167 | 1.024 | 261 | 1.048 | 118 | 1.007 | 46 | 0 | 1 |
| 24 | 179 | 1.022 | 95 | 1.008 | 260 | 1.052 | 47 | 0 | 1 |
| 60 | 181 | 0.992 | 217 | 0.989 | 188 | 0.993 | 48 | 0 | 1 |
| 3 | 154 | 1.011 | 144 | 1.007 | 180 | 1.024 | 49 | 0 | 1 |
| 38 | 142 | 1.016 | 105 | 1.008 | 187 | 1.031 | 50 | 0 | 1 |
| 34 | 146 | 1.005 | 150 | 1.001 | 152 | 1.013 | 51 | 0 | 1 |
| 53 | 151 | 0.994 | 111 | 0.997 | 928 | 0.928 | 52 | 0 | 1 |
| 36 | 143 | 1.013 | 168 | 1.021 | 1936 | 1.801 | 53 | 0 | 1 |
| 42 | 123 | 0.998 | 188 | 0.992 | 123 | 1.006 | 54 | 0 | 1 |
| 33 | 116 | 1.007 | 110 | 1.002 | 135 | 1.002 | 55 | 0 | 1 |
| 5 | 124 | 1.005 | 134 | 0.999 | 140 | 1.004 | 56 | 0 | 1 |
| 66 | 117 | 1.011 | 163 | 1.022 | 113 | 1.004 | 57 | 0 | 1 |
| 57 | 119 | 1.005 | 115 | 1.002 | 140 | 1.009 | 58 | 0 | 1 |
| 48 | 113 | 1.006 | 101 | 1.003 | 271 | 1.062 | 59 | 0 | 1 |
| 39 | 74 | 1.003 | 84 | 1.004 | 108 | 1.002 |  |  |  |
| 32 | 86 | 1.005 | 72 | 1.003 | 107 | 1.013 |  |  |  |
| 64 | 72 | 1.003 | 74 | 1.000 | 100 | 1.009 |  |  |  |
| 28 | 76 | 1.004 | 89 | 1.006 | 86 | 1.005 |  |  |  |
| 31 | 69 | 1.004 | 62 | 1.002 | 118 | 1.013 |  |  |  |
| 29 | 70 | 1.000 | 72 | 1.005 | 119 | 1.015 |  |  |  |
| 27 | 68 | 1.002 | 75 | 1.000 | 200 | 0.992 |  |  |  |
| 1 | 0 | 1.000 | 0 | 1.000 | 0 | 1.000 |  |  |  |

**Table S11.**
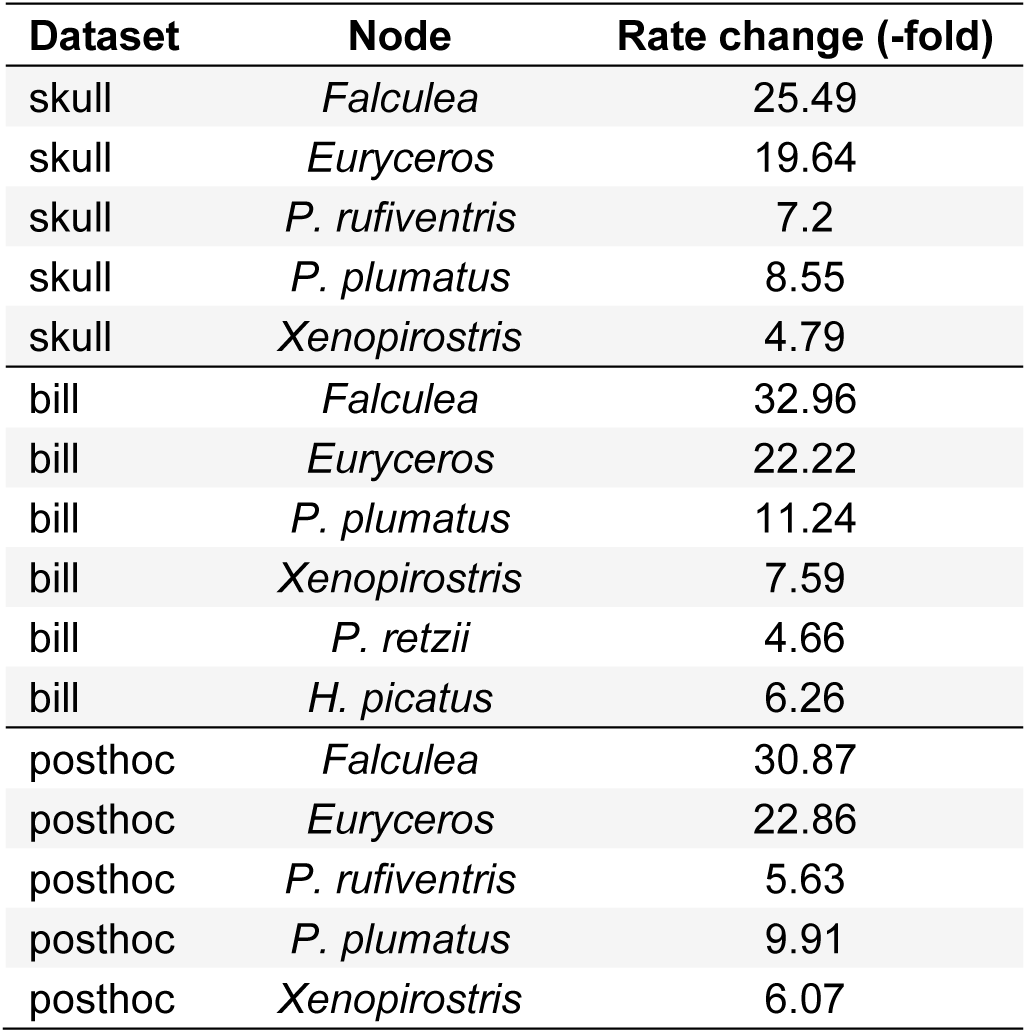
Magnitudes of directional shifts. For each of the highly supported directional shifts reported in Table S10 we calculate the magnitude (fold change) of the shift, as the mean rate is a factor of prior shifts in the ancestors of each species as well as the shift occurring at that branch.

**Table S12.** Significance of net evolutionary rates evaluated using permutation. In Table 3 we reported rate differences for comparisons of the net rate of evolution between clades and foraging classes, as well as the statistical significance of those differences when evaluated using the simulation procedure. However, when significance was instead evaluated using permutation, no differences between clades and few differences between foraging classes were found, with only the very largest rate differences remaining significant. Future work is needed to better understand why these methods produce such different results and appropriate uses of each.

| Module | Malagasy vs. non-Mal | Probe vs. Glean | Probe vs. Sally | Glean vs. Sally |
| --- | --- | --- | --- | --- |
| Whole skull | 0.280 | **0.011** | 0.232 | 0.526 |
| Skull (outliers excluded) | 0.450 | 0.399 | 0.184 | 0.358 |
| Upper bill | 0.099 | **0.012** | 0.453 | 0.174 |
| Lower bill | 0.252 | **0.005** | 0.123 | 0.861 |
| Cranium | 0.694 | 0.380 | 0.808 | 0.435 |
| Feet | 0.186 | 0.974 | 0.304 | 0.160 |
| Feet (outliers excluded) | 0.760 | 0.173 | 0.082 | 0.578 |

